# Time is encoded by methylation changes at clustered CpG sites

**DOI:** 10.1101/2024.12.03.626674

**Authors:** Bracha-Lea Ochana, Daniel Nudelman, Daniel Cohen, Ayelet Peretz, Sheina Piyanzin, Ofer Gal, Amit Horn, Netanel Loyfer, Miri Varshavsky, Ron Raisch, Ilona Shapiro, Yechiel Friedlander, Hagit Hochner, Benjamin Glaser, Yuval Dor, Tommy Kaplan, Ruth Shemer

## Abstract

Age-dependent changes in DNA methylation allow chronological and biological age inference, but the underlying mechanisms remain unclear. Using ultra-deep sequencing of >300 blood samples from healthy individuals, we show that age-dependent DNA methylation changes are regional and occur at multiple adjacent CpG sites, either stochastically or in a coordinated block-like manner. Deep learning analysis of single-molecule patterns in two genomic loci achieved accurate age prediction with a median error of 1.46-1.7 years on held-out human blood samples, dramatically improving current epigenetic clocks. Factors such as gender, BMI, smoking and other measures of biological aging do not affect chronological age inference. Longitudinal 10-year samples revealed that early deviations from epigenetic age are maintained throughout life and subsequent changes faithfully record time. Lastly, the model inferred chronological age from as few as 50 DNA molecules, suggesting that age is encoded by individual cells. Overall, DNA methylation changes in clustered CpG sites illuminate the principles of time measurement by cells and tissues, and facilitate medical and forensic applications.

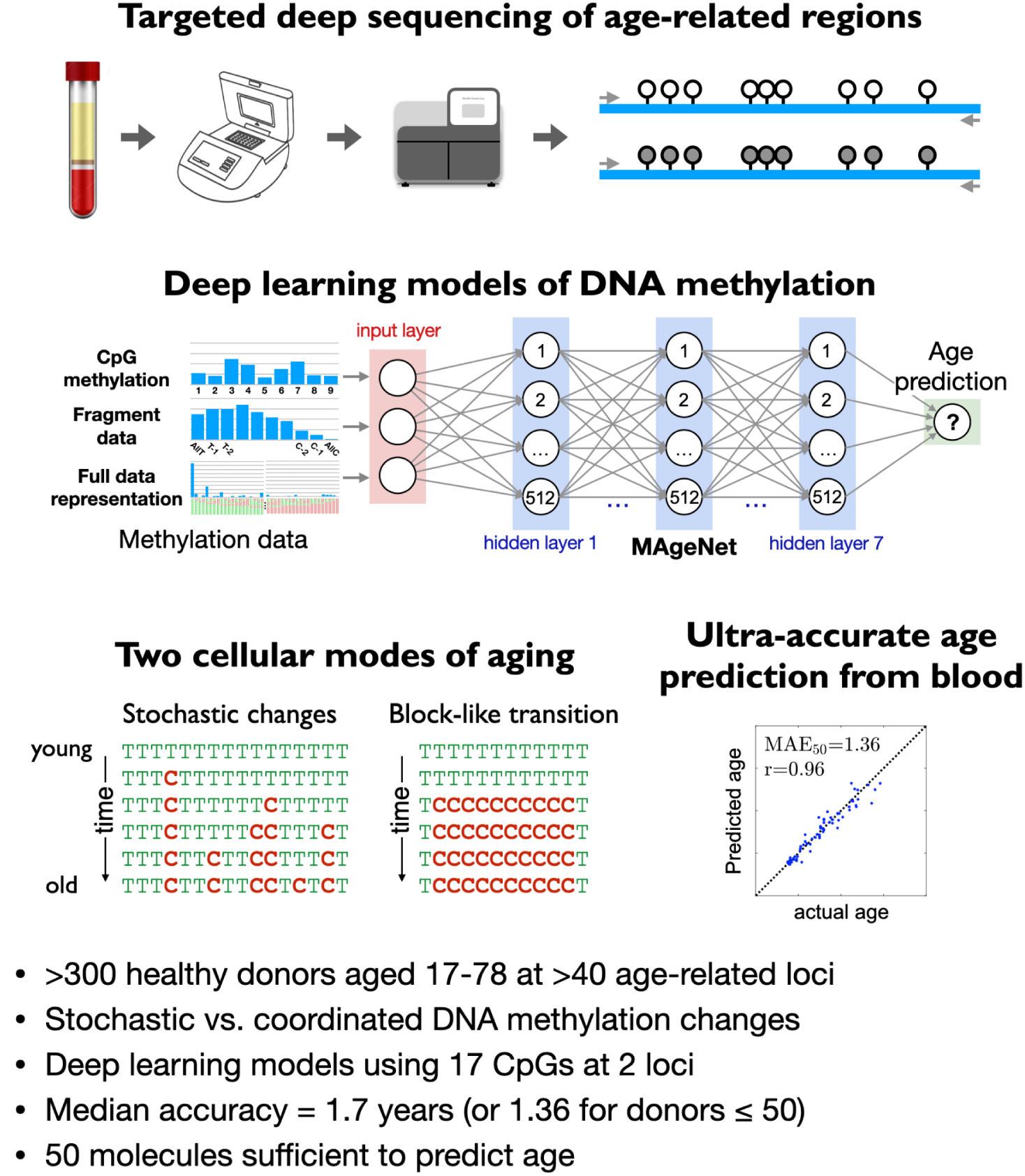

## Introduction

The prediction of chronological and biological age from biological samples offers vast opportunities in clinical diagnostics, monitoring, forensics and aging research^1,2^. Chronological age, defined as the amount of time since birth, correlates strongly with health status; biological age, while harder to define, may provide more accurate information on aging and the propensity for disease^3–5^.

One promising biomarker for age prediction is DNA methylation - the addition of a methyl group to a cytosine in the context of a CpG dinucleotide^5–8^. This epigenetic modification plays a crucial role in various aspects of normal development and disease. During early development, dynamic processes shape the final methylation landscape, which is essential for the specialized functions of various tissues in the developing organism. Once established, these methylation patterns remain stable throughout life and encode the cellular identity of each cell type. DNA methylation regulates and suppresses the expression of silenced genes^9–14^ in a cell-type-specific manner, and indeed the majority of the 28 million CpG sites in the human genome are methylated, whereas gene promoters, enhancers, and CpG islands are often unmethylated, either in all cells or in specific cell types^15–17^. These patterns demonstrate robustness to environmental cues, and present an outstanding similarity across healthy individuals^17^.

However, accumulating evidence has shown that few loci do show dynamic changes in DNA methylation^18^, often associated with aging and diseases such as cancer. Pioneering studies by Horvath and colleagues revealed that a small fraction of CpG sites across the human genome undergo predictable methylation or demethylation with age^5,19^. The methylation patterns of combinations of dozens to hundreds of such sites have been extensively utilized as epigenetic clocks for chronological and biological age prediction^20–22^.

Despite the extensive progress made, current approaches for methylation-based epigenetic age determination suffer from limitations which restrict their accuracy and the biological insight they provide. Epigenetic clocks have been primarily developed using information from Illumina methylation arrays (27K, 450K or EPIC), which measure the average methylation level at a predefined limited set of individual CpGs. As a result, these data cannot detect information embedded in genomic clusters of adjacent CpG sites, and are therefore limited in capturing the full scope of age-related DNA methylation changes. This limitation is significant because DNA methylation does not occur uniformly across the genome; rather, it acts in a regional manner, influencing gene expression, chromatin structure, and regulatory processes in specific regions. Indeed, the biochemistry of DNA methylation dynamics is typically regional, with methylation and demethylation enzymes often acting on multiple adjacent cytosines in a processive concomitant manner^13,23^. Based on this, we hypothesize that age-related changes in DNA methylation occur in a regional manner within clusters of CpG sites, which cannot be measured in a combinatorial way by methylation arrays.

Only few epigenetic clocks based on methylation in adjacent CpGs were proposed^24,25^. Zbieć-Piekarska et al. used targeted pyrosequencing of five genomic CpG sites, and reached a mean absolute error of 3.9 years^26^. TIME-seq, a tagmentation-based approach, reached a mean absolute error of 3.39 years^27^. Yamagishi et al. focused on seven CpG sites located in the promoter of *ELOVL2 and* achieved a mean absolute error of 5.3 years^28^. Finally, the methylation at four age-related regions was shown to predict age in forensic applications, with a median absolute error (MAE) of 5.35 years^29^. Thus, the information embedded in the methylation status of a region with multiple clustered age-responsive sites remains unclear. More recently, single-cell DNA methylation sequencing assays were developed. These approaches capture the methylation status across multiple neighboring CpGs sites, but their sequencing depth is extremely limited, often at 0.1× or below. As a result, only a small fraction of age-related loci are covered across the genome and their general utility in accurately predicting age is limited^30–32^.

A second limitation of current epigenetic clocks concerns data analysis. Most array-based epigenetic clocks are based on linear regression models (e.g. elastic-net^33^), resulting in a MAE of 2.5-5 years^25,34^. Yet, as we and others have shown, methylation in some CpG sites is not linearly correlated with age^18,20,35,36^. Recently, deep neural networks trained on array-based data resulted in an MAE of ∼2.2-2.7 years, using ∼1000 CpGs^37^. We have recently reported GP-age, a non-linear cohort-based computational algorithm that further improves on current clocks, resulting in a MAE of 1.89-2.1 years based on 30-80 CpGs selected from the 450K and EPIC arrays^18^. Finally, the cost and turnaround times of array-based measurements are typically high, limiting utility relative to targeted analysis of few informative loci.

Here we present a novel framework for methylation-based chronological age determination that integrates targeted DNA methylation sequencing of selected loci with deep neural network analysis. We examined 45 CpG sites previously reported to be age-responsive, and used multiplex targeted-PCR followed by ultra-deep sequencing to determine the methylation status of these CpG sites, along with multiple adjacent CpGs within the same region, using genomic DNA from blood samples obtained from 300 healthy donors across three independet cohorts. This allowed us to explore age-related DNA methylation changes that occur in a regional manner within clusters of CpG sites, revealing that some regions change stochastically and others in a block-like coordinated manner. We then developed a novel framework for methylation-based chronological age prediction, integrating single-molecule combinatorial patterns across multiple methylation sites. We trained a fully connected deep neural network, resulting with a robust epigenetic clock that obtains a median accuracy of 1.46-1.7 years on held-out samples, regardless of environmental factors. We explored the impact of environmental factors. We further explore the minimum number of cells required to encode elapsed time, and discuss potential applications in forensics and in aging and rejuvenation research.

## Results

### The methylation neighborhood of age-related CpGs

To explore the nature of CpGs sites surrounding age-related CpGs, we began by focusing on data from Illumina 450K and EPIC DNA methylation arrays, identifying CpG sites highly correlated with age, and examining the methylation of neighboring sites. This allowed us to study how likely it is to establish a computational analysis of multiple neighboring CpGs, sequenced together, and to assess methylation dynamics of CpGs in the vicinity of age-related sites. Utilizing a published dataset of 11,910 blood methylomes of healthy donors aged 0-103 years^18^, we focused on CpGs that exhibit strong correlation with age (absolute Spearman correlation coefficient rho≥0.4) and show a large absolute change of DNA methylation levels during adulthood (a change of ≥ 20 percent points between ages 20 and 80). These thresholds identified a total of 2,374 age-related CpG sites, for which we examined the presence of neighboring CpG sites (up to 450 bp, to fit within a single amplicon), and their correlation with age. Remarkably, nearly 70% of age-related sites have neighboring CpGs (Fig. 1A). Incidentally, 10% of these neighboring sites are included in the methylation array design, allowing us to calculate their correlation with age. CpGs within 50 bp from the top 2,374 age-correlated CpGs are strongly correlated with chronological age (Spearman |ρ|>0.35, Fig. 1B), suggesting that age-related methylation changes often occur across multiple neighboring sites, rather than at individual positions.

**Figure 1:**
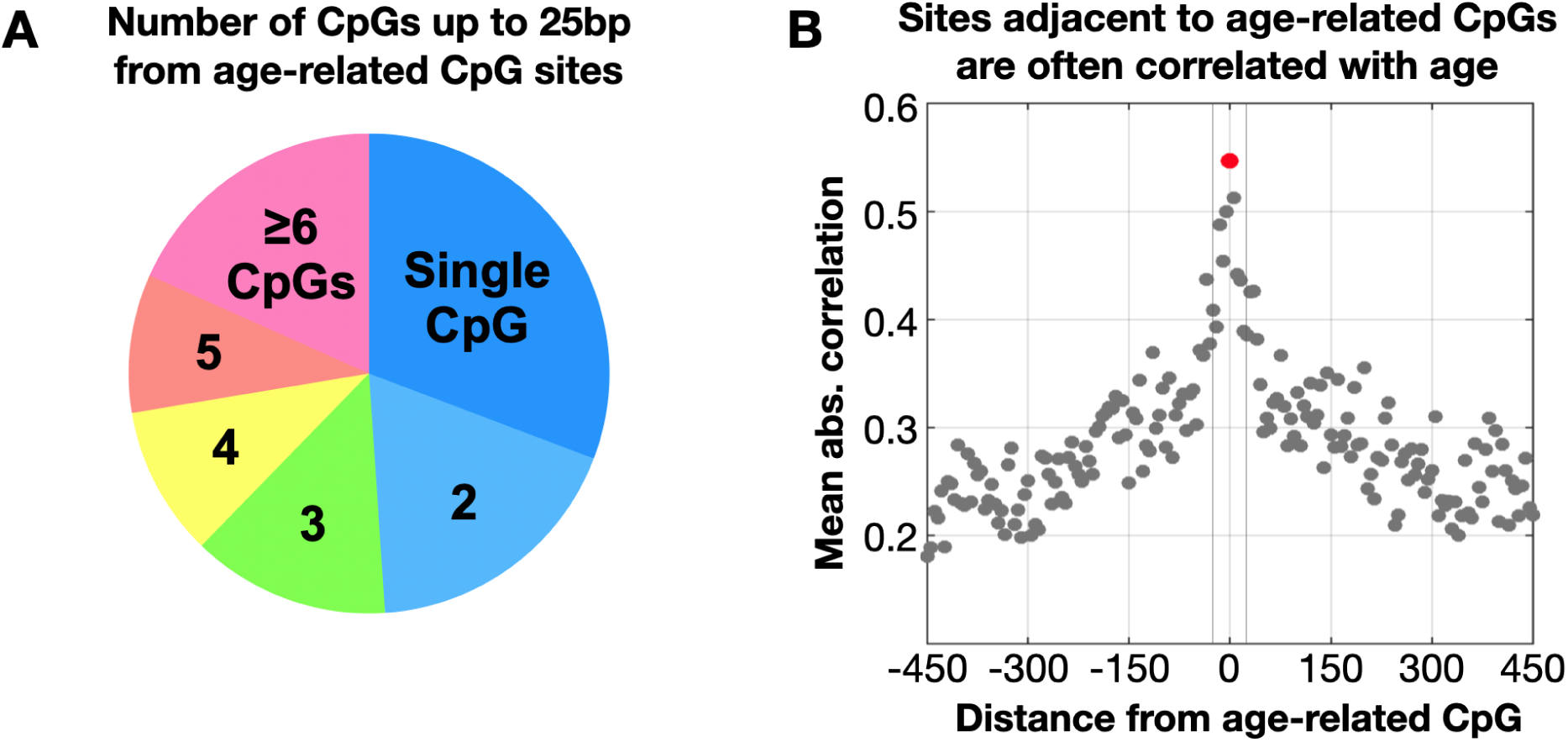
450K/EPIC age-associated DNA methylation sites are often surrounded by additional CpGs correlated with age. **(A)** Top 2,374 age-correlated sites were identified using 11,910 blood DNA samples from Varshavsky et al.^18^. Of these, only 31% are single (blue), whereas most 450K/EPIC age-related sites are surrounded by multiple CpGs (up to 25bp away), which are typically not measured using DNA methylation arrays. **(B)** The average correlation between DNA methylation and age is shown for the top 2,374 sites (red dot, center), as well as neighboring CpG sites that are present on the methylation array (gray dots, grouped by relative distance).

Consequently, we hypothesized that a targeted bisulfite-PCR approach followed by ultra-deep next-generation sequencing, could shed light on age-related methylation dynamics by measuring the combinatorial patterns of multiple CpGs in thousands of DNA fragments at a single-molecule resolution (Fig. 2A).

**Figure 2:**
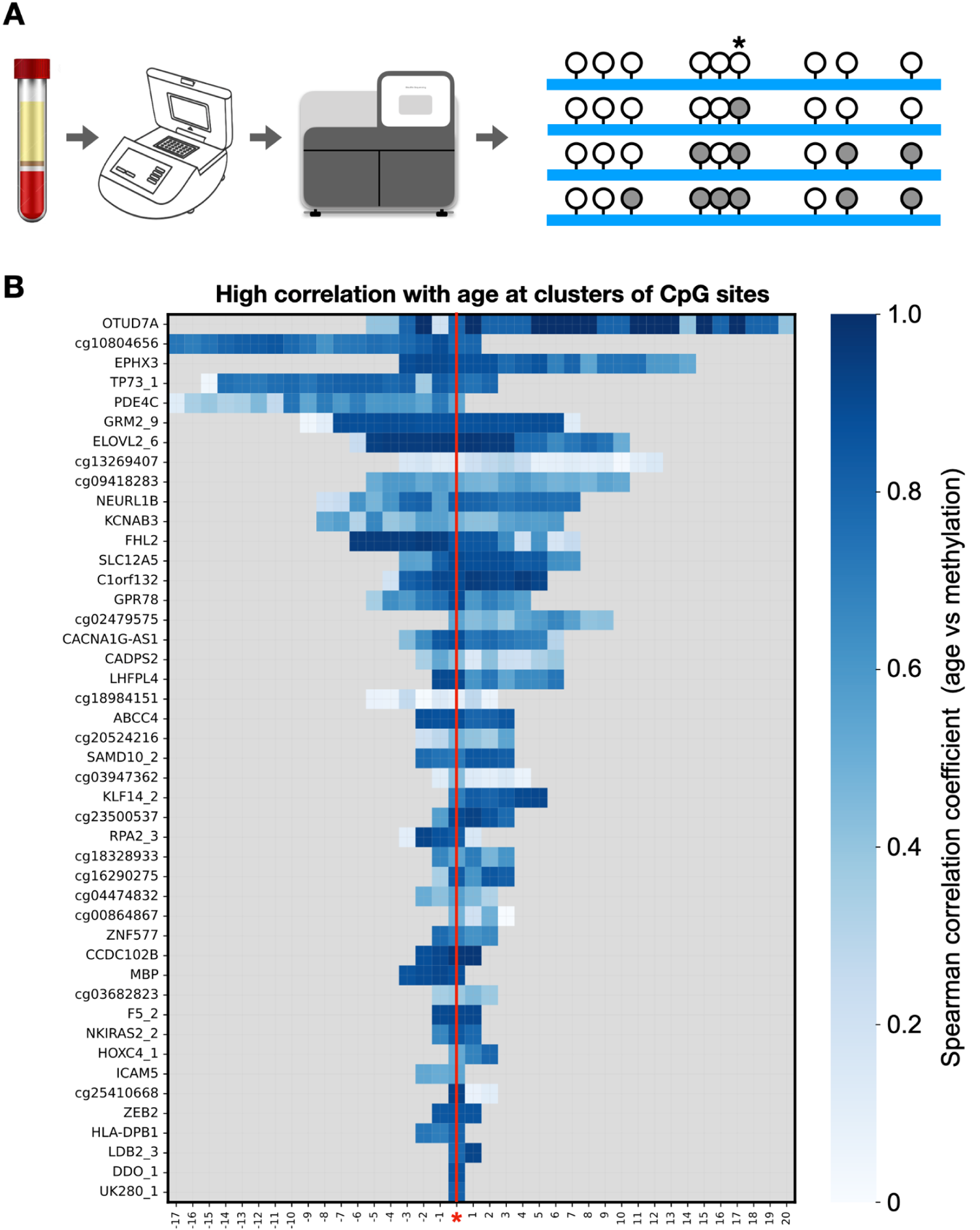
Clusters of age-related changes. **(A)** Schematic of targeted PCR-sequencing following bisulfite conversion, facilitating concurrent mapping of multiple neighboring CpG sites at a depth >5,000×. Red and green, methylated and unmethylated CpGs; asterisk, CpG present in array. **(B)** For each amplicon (row) we plot the Spearman correlation coefficient for each CpG, showing high concordance between age and DNA methylation levels across multiple, clustered, CpG sites. Amplicons are aligned by DNA methylation array CpG.

### Development of age-responsive multi-CpG markers and a cohort of healthy donors

To identify clusters of CpGs whose methylation strongly correlates with age, we considered the most correlated sites from the published dataset of 11,910 blood methylomes, as well as additional genomic regions previously associated with age^18,20,21,34^. We expanded each of these array-based CpG sites to include nearby CpG sites, and designed targeted PCR primers to co-amplify and sequence 45 target regions following bisulfite conversion. The amplicons were on an average 134bp long, and had 8.5 CpGs per amplicon, and represent a mixture of genomic regions that either gain or lose methylation with age, including gene promoters, introns, CpG islands, polycomb CpG islands, flanking regions, distal enhancers and more (Table S1).

We then collected blood samples from 296 self-declared healthy donors, aged 17 to 78, extracted DNA, and treated with bisulfite^38^. This was followed by multiplexed PCR amplification of over ∼2000 genome equivalents (10 ng), and sequencing at an average depth of 12,839 fragments per amplicon (Table S1). Samples were then divided into training set samples (n=205, 42 of which marked as validation samples for hyperparameter tuning), and held-out test samples (n=91), after stratifying by age (Fig. S1). The three sets show similar age distributions and are balanced for gender. Additional data provided by the donors included weight, height, smoking status, smoking years, and brief medical history (Table S2). Figure 2B shows the absolute Spearman correlation between DNA methylation and age, for each individual CpG site we measured. Each row shows one such genomic region (amplicon), centered by the original age-related CpG site from the methylation array design, with surrounding CpGs spanning to the right and left. Indeed, at most amplicons, we observe a number of multiple age-related CpG sites are clustered in proximity (Fig. 2B, Table S3).

### Clustered, stochastic, non-linear methylation changes at the ELOVL2 locus

We begin by examining how blood DNA methylation changes with age, across a set of adjacent CpG sites in one particularly informative locus. In Fig. 3, we illustrate the average methylation of 17 sites at the 154bp-long ELOVL2 amplicon (chr6:11044843-11044997, hg19), across 205 training samples aged 17-78. As previously showed, CpG #7 in our amplicon (cg16867657) is highly correlated with age. Yet, it is a part of nine CpG sites (#2-10), located within a small region of 57bp, that show a dramatic consistent accumulation of methylated molecules with advanced age. Remarkably, despite their physical proximity (CpGs #5-7 are directly adjacent, others are within few bases), these age-responsive CpG sites all show different dynamics throughout life - they show a range of baseline methylation levels (CpG #2 vs #4, or #8 vs #9), as well as differences in their annual rate of change. Additionally, these sites are flanked by a CpG site (#1) that shows no DNA methylation changes whatsoever; as well as a group of seven CpGs (#11-17) that show a small but consistent rate of annual DNA methylation gain (Figs 3, Table S3).

**Figure 3:**
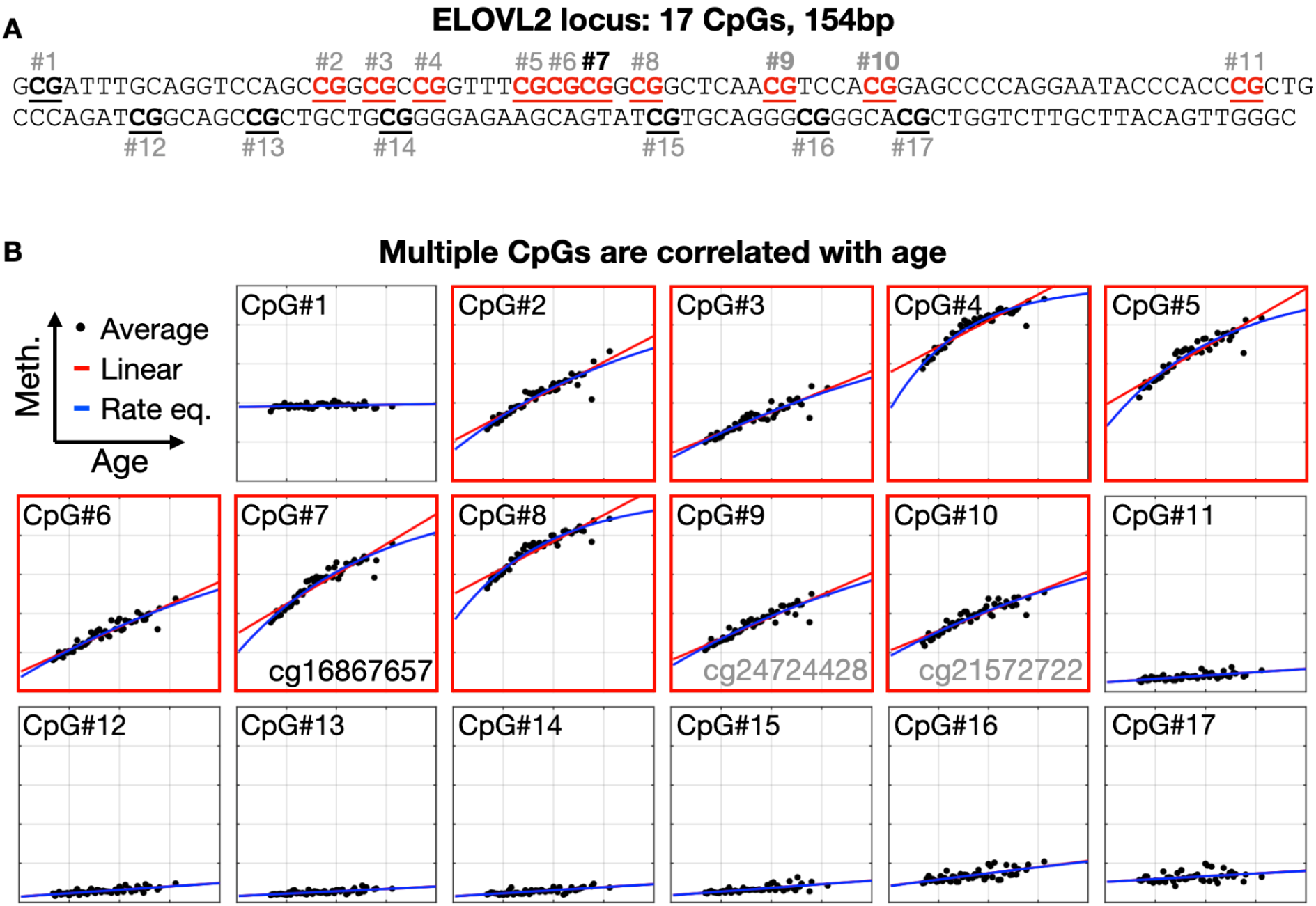
Clustered, non-linear age-related methylation changes. **(A)** Shown are 17 CpG sites from the ELOVL2 amplicon (chr6:11044843-11044997, hg19). Each dot represents the average DNA methylation from deeply sequenced blood DNA, from a single donor. X-axis: chronological age on a 0-100 scale; Y-axis: methylation on a 0-100% scale. Marked in red are CpG sites strongly associated with age, with absolute Spearman correlation ≥ 0.8 and methylation range ≥ 20 percentage points. Data points are fitted using a linear model (red line) or a first-order rate equation (blue line). **(B)** DNA sequence at the ELOVL2 amplicon. Age-associated sites are highlighted in red. ELOVL2 CpG #7 (cg16867657) is marked.

Intriguingly, the commonly used linear models, that assume a constant change in methylation levels during adulthood^39–42^, provide a rather poor fit for most age-specific changes. Conversely, we show that a simple rate equation, by which a fixed percent of unmethylated molecules changes each year, offers a better fit for most CpG sites (Fig. 3). Specifically for CpG#7 (cg16867657), used by many epigenetic clocks, the non-linear fit is significantly more accurate than the linear fit, with RMSEs of 2.8% vs 3.4%, respectively (p≤3.6e-5). Importantly, the rate equation model offers a mean prediction error of 3.2 years (based on a single CpG site), compared to 4.2 years when using the linear model. Similar principles were observed for additional CpGs at multiple regions, including regions that demethylate with age (Fig. S2).

Overall, age-correlated CpGs are often located in clusters, and the sites measured by methylation arrays are often not the ones to change the most (Fig. S3). Suggesting that neighboring sites, and especially the combinations of multiple sites, could offer an improved age prediction from methylation.

### Stochastic vs. block-like methylation changes at neighboring CpG sites

To better understand the dependencies between methylation of neighboring CpGs, we focus on the methylation patterns of individual single molecules. Each sequenced read was analyzed, and the binary patterns of covered CpGs was recorded. We then examined the frequency of each possible binary pattern at donors of different ages.

Indeed, we identified two very distinct modes of age-related methylation changes that are undistinguished when examining data from DNA methylation arrays. Some genomic regions show stochastic, position-independent methylation changes, by which each individual CpG is randomly changed, independently of changes in neighboring sites (Fig. 4A, top). Conversely, other genomic regions seem to show a block-like coordinated change of multiple neighboring CpG sites (Fig. 4A, bottom), by which all adjacent sites exhibit the same methylation level, and resulting with a mixture of fully methylated and fully unmethylated molecules.

**Figure 4:**
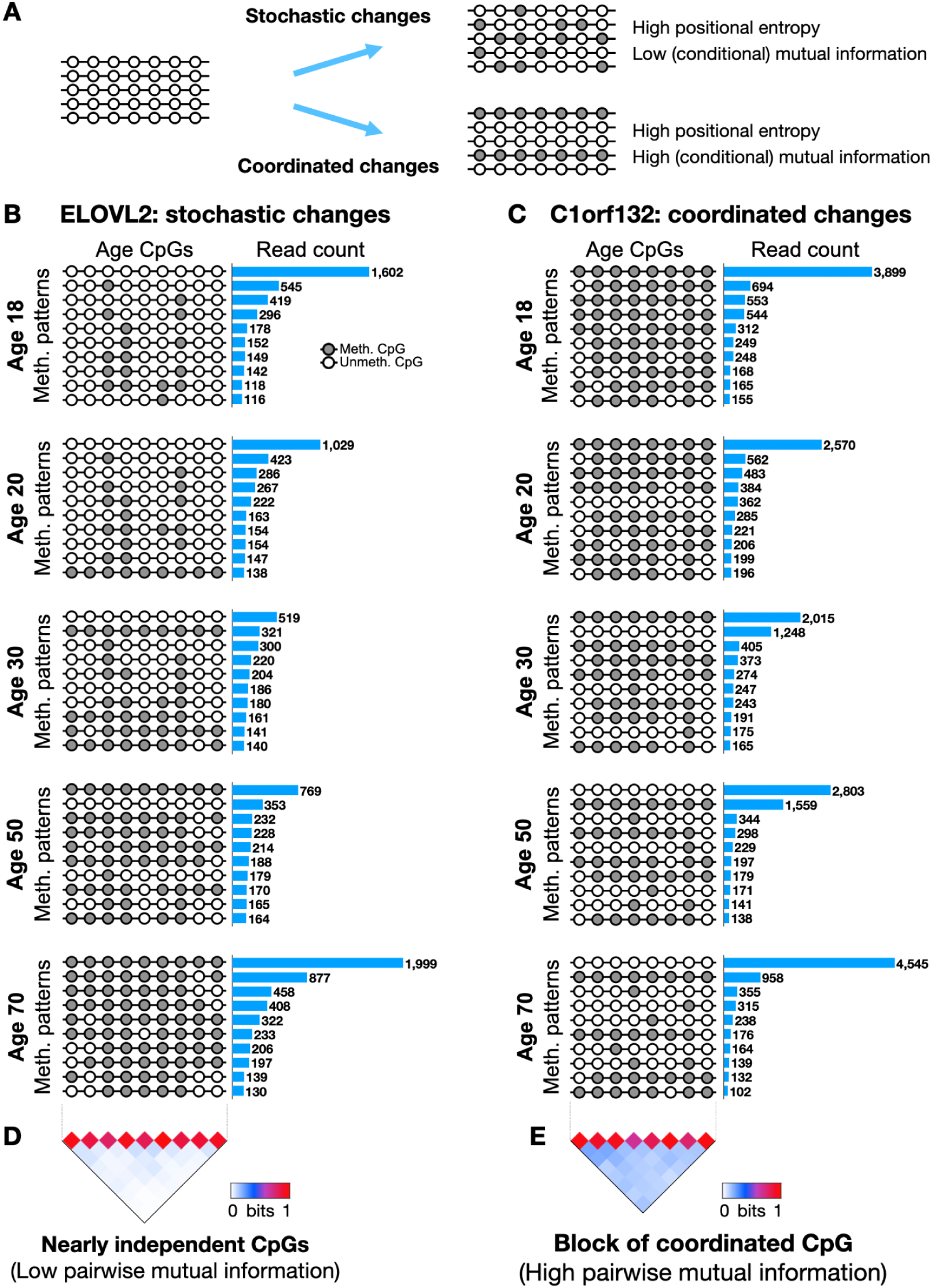
Stochastic vs. block-like age-related methylation changes at neighboring CpG sites. **(A)** We propose a model by which an unmethylated region could gain methylation by stochastic accumulation of individual changes, or by block-like concordant changes across multiple CpG sites. **(B)** The observed frequency of multiple binary patterns across nine CpGs sites, at the ELOVL2 locus, at five ages (18, 20, 30, 50 and 70). A strong lifetime gain of methylation is observed, by which the fully unmethylated molecule (all white) becomes less frequent as more and more CpG sites are randomly methylated, until this region is mostly methylated at older ages. **(B)** Same as (A), for the C1orf132 locus. Here, the CpG sites are strongly related, and the fully methylated pattern is replaced over the years by the fully unmethylated pattern. **(D)** The ELOVL2 locus is characterized by highly variable CpGs (with high entropy) that are largely independent of each other (low pairwise mutual information). (E) Conversely, C1orf132 is characterized by strong pairwise coordination.

At the ELOVL2 locus (Fig. 4B), young donors are characterized by unmethylated DNA fragments across all nine age-responsive sites (CpGs #2-10). This pattern gradually gives way to an ensemble of “dotted” molecules with mixed methylation, until fullly methylated molecules become the most abundant pattern at older donors (Fig. 4B).

To quantify how coordinated each pair of CpGs is, while considering individual age-related changes, we devised a computational score based on conditional mutual information. This score measures, for donors of every age, how much of the uncertainty of one CpG site is reduced by knowing the methylation state of the other site (in bits). We then average across all ages to quantify the overall mutual information between two CpGs.

For highly variable CpGs, we observe high entropy for each individual site, but near-zero pairwise mutual entropy, suggest that the sites are mostly independent. Conversely, high pairwise values suggest that the two CpGs change concordantly, in a coordinated way. To account for age-related dynamics, the mutual information score we calculated was estimated for each age independently, then weighed and summed across all ages (see Methods). As shown in Figure 4D, the nine CpGs at the ELOVL2 locus are nearly independent of each other, with near-zero pairwise mutual information.

Intriguingly, we also observed a second mode of age-related methylation changes, characterized by coordinated demethylation of multiple neighboring CpGs. Figure 4C depicts the abundance of patterns across a neighboring set of eight age-related CpG sites at the C1orf132 locus. Unlike the stochastic changes noted in ELOVL2, age-related methylation changes in this locus occur in a block-like manner, by which nearly all CpGs are either methylated or unmethylated. These block-like concordant changes across multiple neighboring CpGs is typical of cell-type-specific differentially methylated regions, as previously reported by us and others^17,23,43^, but not in the context of cellular aging.

Indeed, a pairwise analysis of the C1orf132 amplicon using conditional mutual information, identified a coordinated block of eight CpGs (Fig. 4). These two archetypical modes of change were observed in additional age-responsive amplicons that we tested, including TP73 and CCDC102B (stochastic) and FHL2, SPAG9 and GRM2 (block-like) (Figs. S4). These two principles are further visualized in Fig. S5, showing a gradual stochastic accumulation of methylation, shifting from the fully unmethylated to the fully methylated pattern (or vice versa), for some regions; alternatively, other block-like regions directly switch from one pattern to the other, without going through the interim, mixed, patterns.

Recently, Tong et al examined the stochastic processes that underlie epigenetic clocks^44^. Importantly, their analysis and simulations were based on 450K and EPIC arrays, limited to average methylation values at individual sites, and therefore cannot accommodate the different principles of coordinated and independent changes described here. In contrast, the 45 regions in our study show a combination of DNA methylation changes that accumulate as we age at a single molecule level, either stochastically (at individual CpG sites), or in a coordinated manner (across a set of CpG sites).

Moreover, our results warrant biochemical examination of the mechanism of DNA methylation change with aging, including analysis of chromatin accessibility and processivity by methylation enzymes e.g. TET and DNA methyltransferases.

### MAgeNet, a deep neural network for chronological age prediction

Next we devised an epigenetic clock, based on multiplexed PCR followed by sequencing, to infer chronological age. Unlike previous approaches that predicted age from average methylation levels at individual CpGs, we wished to integrate the combinatorial methylation patterns of multiple CpGs sites at individual DNA molecules, in thousands of sequence reads from few age-related regions. For accurate and robust predictions, we designed three complementary representations for each sample, reflecting different degrees of abstraction and processing. For a locus with K age-related CpGs, the first representation holds the average methylation at each individual CpG; the second representation contains the abundance of fully unmethylated reads, of reads methylated at exactly one CpG, at two, three, and so on, (a total of K+1 features, from 0 through K); the third representation contains the abundance of each possible combinatorial pattern across the K CpGs, to a total of 2^K^ possible options (Fig. 5A).

**Figure 5:**
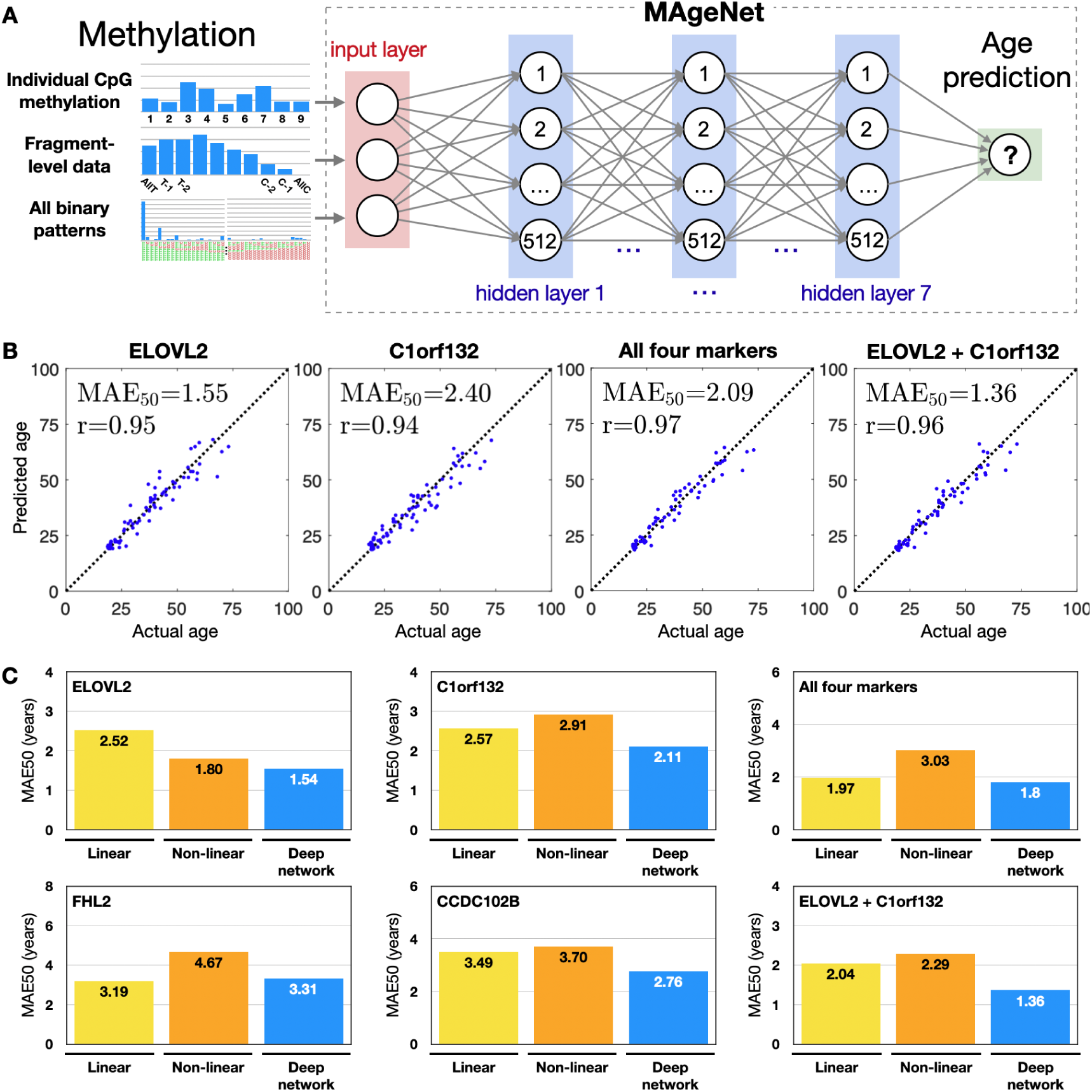
MAgeNet outperforms other linear and non-linear regression models for age prediction from DNA methylation. (A) A deep neural network for age prediction from fragment-level targeted DNA methylation data. Following targeted sequencing, each library was represented using individual CpG methylation levels, the abundance of fragment-level counts of methylated CpGs (e.g. “all-but-one”), and a combination of all binary patterns. These were the input of a 7-layer fully connected deep network, with non-linear ReLU activation functions and 256 or 512 neurons per layer. (B) Deep learning predictions (y-axis) vs actual age (x-axis) are shown for the ELOVL2 locus, for C1orf132, for all four markers (including FHL2, and CCDC102B), and for a combined two-marker model. The latter model achieved a median abs. error of 1.7 years on held-out test-set donors (or 1.36 years for donors aged 50 or less). (C) A comparison of median errors (Y-axis) for linear, non-linear, and deep learning models. The most accurate predictions are typically obtained by deep learning models (blue), rather than the commonly used linear model (elastic-net, green), or a generalized non-linear model (orange). Overall, the most accurate predictions were achieved by deep learning models. ELOVL2 achieved a median error of 1.8 years (1.54 for donors ≤ 50), or a combination of ELOVL2 and C1orf132, with a median error of 1.7 years (1.36 years below 50), using multiplexed targeted PCR sequencing data.

Overall, of the 45 genomic regions we measured, 16 genomic regions showed a dramatic change in at least three consecutive CpG sites, defined by an absolute Spearman correlation of 0.8, as well as an absolute change in methylation of 20 percentage points during adulthood (Methods, Table S3). We then trained various models, including the linear elastic net regression model, the non-linear GAM regression model^45^, and deep learning (fully connected neural networks)^46,47^, using the three different representations of the data. This allowed us to quantify the importance of different representations and the accuracy obtained for different loci by each model. For improved robustness, and to account for the varying number of sequenced reads per sample, the sequencing data (for both training and test-set samples) was augmented by generating 128 random subsets of 8,192 reads each, by sampling (with replacement) from the original data. This also accounts for the variable number of sequenced reads from each donor or locus. A principal component analysis of these high-dimensional data revealed that 92% of the variance could be explained by a single dimension, which is also highly indicative of age (Fig. S6).

We designed MAgeNet, a deep fully connected neural network for chronological age prediction from targeted PCR-based DNA methylation sequencing from blood (Fig. 5A). Hyper-parameters were selected using a grid search and L1 loss (on the validation set), and the optimal model for each marker was then retrained on all training data (see Methods). We also trained regression models for each amplicon and tested the models on the held-out test-set samples. Four genomic regions: ELOVL2, C1orf132, FHL2, and CCDC102B, showed comparable prediction accuracy, with MAE≤4 years, and root mean square error (RMSE) below 7 years (Table S4). For forensic applications, we also calculated the MAE and RMSE scores for donors aged 50 or younger (MAE50, RMSE50, Figs. 5, S7, Tables S5-S6).

### Ultra-accurate age prediction from blood DNA methylation

Strikingly, we found that PCR-based targeted bisulfite sequencing, capturing the combinatorial patterns of multiple age-related neighboring CpGs from just one genomic locus, outperforms all known epigenetic clocks. For example, a deep learning model trained on the ELOVL2 locus solely, composed of nine age-related CpGs, achieves a MAE of 1.8 years for held-out test samples; or 1.54 years for donors below 50 (MAE50, Figs. 5, S7). A model based on eight CpGs at the amplicon at C1orf132 achieves a MAE50 of 2.1 years; the nine-CpG amplicon near FHL2 yields a MAE50 of 3.3 years, and a model based on four CpGs near CCDC102B presents a MAE50 of 2.8 years.

The models we trained also allow us to compare, for each amplicon and in an unbiased way, how different representations and different models affect prediction accuracy. For the stochastic ELOVL2, the full combinatorial representation was as accurate as the simpler representation based on how many CpGs are methylated, in each individual read, and both representations outperformed the common representation of beta values (average methylation levels) at individual CpGs. Importantly, regardless of data representation, the two non-linear models - generalized additive models (GAM) and deep neural networks - outperform linear regression models.

Next we combined multiple amplicons to increase the accuracy and robustness of age prediction. We merged the feature-based representation of each marker and trained a larger network. Indeed, a joint end-to-end model of two loci (ELOVL2 and C1orf132) outperformed all single-locus models, and achieved a median absolute error of 1.7 years across the test set samples. Importantly, the model’s median accuracy for donors 17-50 years old was 1.36 years, and 0.9 years for test-set donors between 17 and 35 years old (Figs. 5, S7), thus offering state-of-the-art accuracy for various applications in forensics, medicine, and aging research.

### Minimum number of cells needed for age inference

We next examined the minimal number of cells required to accurately predict chronological age, as information on this matter may shed light on principles of aging, and have practical implications in forensics, where the amount of available material is often very limited. For this, we took two complementary approaches. First, we sub-sampled our sequenced PCR libraries to simulate lower library complexity and sequencing depth. For example, when simulating 100 cells, we randomly sampled 100 reads for each locus (for each donor), effectively reducing the average depth of sequencing for the whole dataset from 12,839× to 100×. We then repeated the age prediction pipeline described above. Overall, we applied this procedure 100 times with n=10, 20, 50, 100, 200, 500, 1000, 2000, 5000 and 10000 sampled reads. As Figure 6A shows, 500 DNA molecules per region are sufficient for highly accurate age prediction, with a median accuracy (MAE50) of 1.53 years (±0.2). Even DNA equivalent to 20-50 cells was sufficient to predict age with a median absolute error of 3-4 years. These results reflect upon the ability to predict age from single-cell DNA methylation data, where the sequencing depth per cell is extremely low (at ∼0.1×), and the overall coverage per locus is low^30,31^.

**Figure 6:**
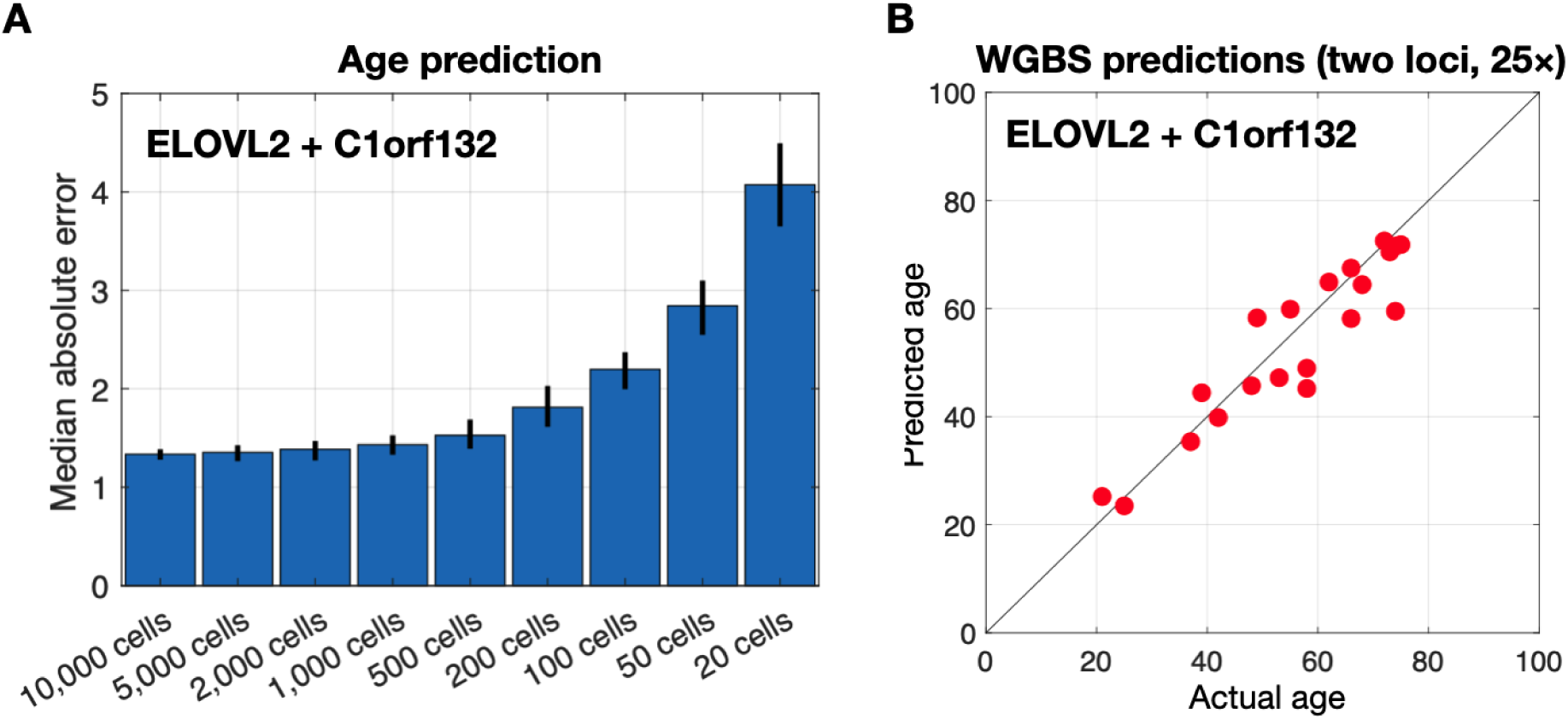
Age predictions are robust to environmental and physical characteristics. **(A)** Comparison of donor sub-groups shows no bias to age prediction (Y-axis) introduced by sex, body-mass index (BMI) and smoking status. **(B)** A comparison of predicted epigenetic age (X-axis) vs predicted biological age (Y-axis), using blood samples from the Jerusalem Perinatal Study (JPS), shows no effect of biological age on the predicted epigenetic age. Red lines mark the average chronological age of the group. **(C)** Longitudinal analysis of blood samples from the Jerusalem Perinatal Study (JPS) cohort revealed a median error of 1.73 years for donors aged 30-33 (red dots). Analysis of blood taken from the same patients 10 years later (blue dots) showed a relative median error of 1.38 years, suggesting that deviations in epigenetic age prediction are due to earlier life events or genetics, and that methylation changes at the ELOVL2 and c1orf132 amplicons accurately record the passage of time.

**Figure 6:**
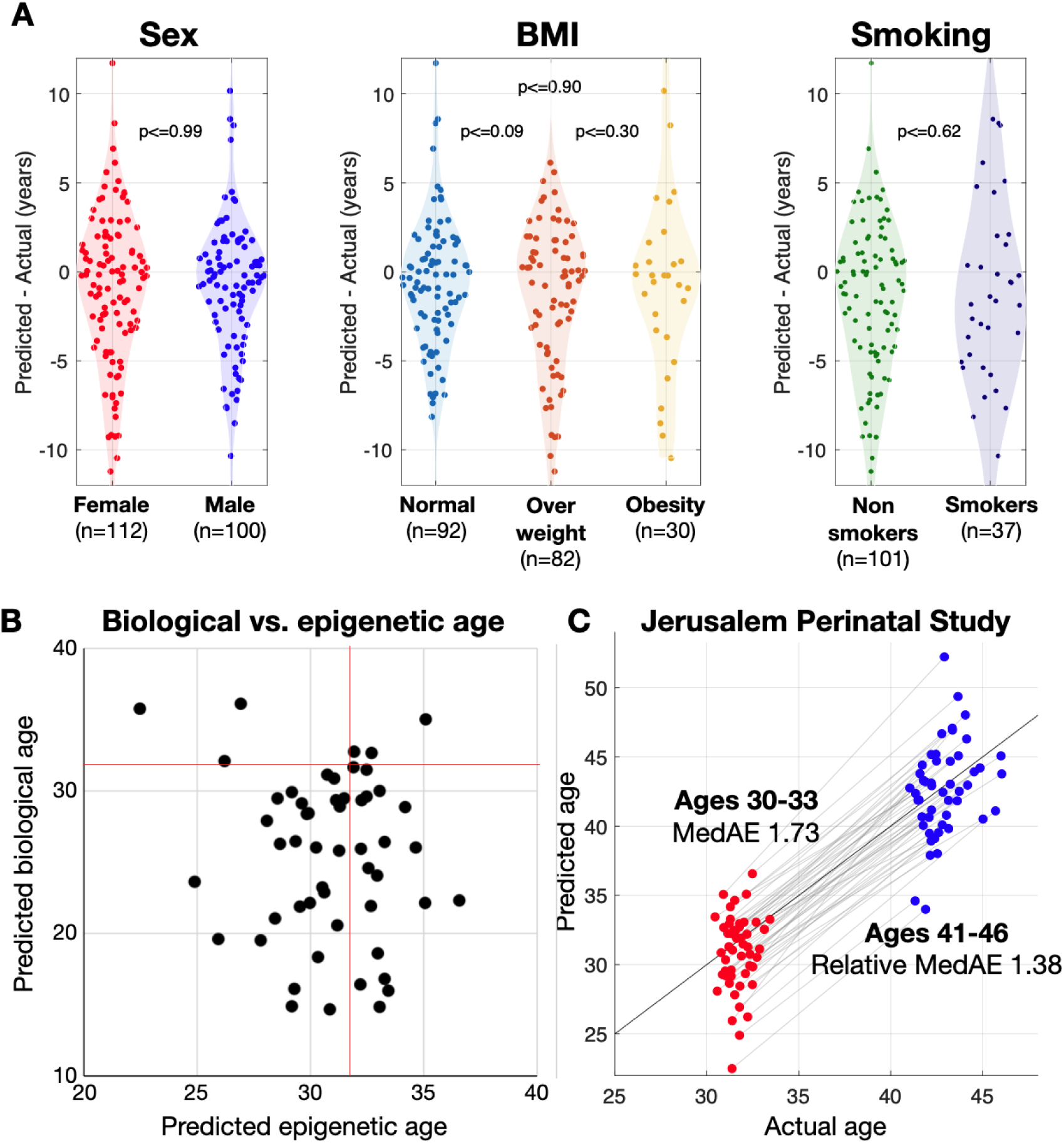
Age predictions are robust to environmental and physical characteristics and require few cells. **(A)** Sampling analysis shows the expected median error (y-axis) from increasingly smaller PCR libraries, demonstrating accurate age predictions from as little as 200-500 sequenced PCR products. **(B)** Age prediction from whole-genome bisulfite sequencing^17^ shows accurate predictions at an effective depth of 25× in two loci, suggesting that age could be inferred from fewer than 50 blood cells.

Second, we applied MAgeNet to 23 whole-genome bisulfite sequencing samples we recently published^17^, containing methylation data of genomic DNA from white blood cells from healthy adults (mean age 57 years). Focusing on reads that fully cover the ELOVL2 and C1orf132 loci (∼25×), we applied MAgeNet and achieved a median absolute accuracy of 3.58 years (Figs. 6B, S8). These results provide an independent validation for the performance of our clock, and support the idea that a small number of cells may suffice for age prediction.

### Age-dependent vs cell-composition DNA methylation changes

The findings described above are based on methylation of DNA from whole blood, and are potentially affected by differential cell type composition (which is known to change with age)^48,49^. To assess the performance of our methylation markers in predicting age from pure cell types, we collected DNA from eight healthy donors aged 25 to 63 and FACS-isolated neutrophils, monocytes, B cells and T cells. We extracted DNA, treated with bisulfite, PCR-amplified and sequenced to an average depth of 12,303× (Fig. S9). Interestingly, we found that some age-related methylation changes reflect slow changes in the cellular composition of blood DNA, rather than a gain or loss of methylation^18,50^. For example, the C1orf132 locus is losing methylation in some cell types, but remains hypermethylated in T cells (Fig. S10).

Importantly, the nine age-correlated CpGs in the ELOVL2 locus showed no significant bias across blood cell types, suggesting that methylation changes in this locus are primarily driven by age. These results suggest an intriguing interpretation of the deep neural network, by which the integration of ELOVL2, a cell-type-independent marker, with C1orf132, a cell-type-dependent marker, improves prediction accuracy by integrating a wider spectrum of age-related phenomena.

### Chronological age prediction is not influenced by measures of biological aging

We next turned to examine whether individual characteristics affect the prediction accuracy of our targeted epigenetic clock. Higher BMI was previously associated with epigenetic age acceleration^51,52^. Smoking-related methylation alterations were also reported^53^, although the effect of smoking on age-responsive methylation is unclear. Sex differences were also shown to affect epigenetic clocks^54^. To explore these factors we divided donor samples based on these criteria and determined the accuracy of prediction in each group. We found no effect of smoking status, BMI and sex on the accuracy of age prediction (Figure 6, Table S7). These findings suggest that the two-loci chronological age predictor we trained is robust to environmental and hormonal cues.

To validate these findings, we turned to another independent cohort. The Jerusalem Perinatal Study (JPS) monitors thousands of individuals born in Jerusalem between 1964 and 1976^55,56^. We analyzed blood samples from 52 donors, taken 10 years apart, at the ages of ∼32 and ∼42, and processed the samples as described above. We then applied the two-loci (ELOVL2, C1orf132) epigenetic age model and predicted chronological age. Overall, the median accuracy of MAgeNet was 1.73 years, providing an independent validation to the performance of the algorithm.

Alongside the chronological age of the JPS donors, we analyzed various biological measurements including blood glucose, total cholesterol, triglycerides, BMI, waist circumference, and diastolic and systolic blood pressure. For the initial time point (age 32), additional measurements were available, including blood urea nitrogen (BUN), creatinine, uric acid, C-reactive protein (CRP), alkaline phosphatase (ALP), albumin and an estimate of biological age (BA), as predicted based on these biomarkers^57^. We found that none of the biological measurements affected chronological age prediction (Fig. 6), with one exception: for ∼32 years old donors, higher triglycerides levels in the blood seem to affect the epigenetic age prediction error (Spearman −0.3, FDR ≤ 0.017, Table S8, Fig. S11). We speculate that methylation of ELOVL2 may affect the function of its gene product, a fatty acid elongase. Nevertheless, this effect was not observed for the group of ∼42 year old participants, further strengthening our claim of the clock robustness and independence from biological age.

### Consistent 10-year longitudinal predictions

We compared the deviations of the clock predictions from chronological age for the two time points in the JPS cohort. At 32, the MAgeNet predictions obtained a median accuracy of 1.73 years, compared to median accuracy of 2.2 years, 10 years later. We therefore focused on each individual donor, and compared if the two deviations were coordinated. Indeed, the difference between the two predicted ages was highly correlated with the chronological difference between tests, with a relative median error of 1.38 years (Fig. 6C). In other words, a pre-existing deviation between actual and predicted epigenetic age is likely to be carried over to the future, and current deviations could indicate early life events or genetic factors that affected the clock in the past, after which the passage of time was faithfully recorded.

## Discussion

Most methylation-based epigenetic clocks were developed to reflect chronological as well as biological age, such that deviations from chronological age are interpreted as a reflection of accelerated or decelerated aging. We aimed to target the molecular mechanisms that encode purely chronological age, to better understand the underlying biology of how elapsed time is encoded in cells and to provide tools for research and forensic applications. The approach that we developed is based on two principles. First, targeted PCR-sequencing of selected age-responsive loci, to assess the methylation status of multiple neighboring CpGs; Second, deep learning based on fully connected neural networks utilize non-linear activation functions at each artificial neuron. MAgeNet, the resulting algorithm, offers a compelling alternative to DNA methylation arrays, with a dramatic improvement in the accuracy of chronological age prediction at reduced cost and a faster turnaround time. Performance was assessed using held-out samples from our collection of 300 blood samples, and further validated using samples from two independent cohorts – a 10-year longitudinal analysis of 52 donors from the Jerusalem Perinatal Study, as well as 23 donors subjected to WGBS of genomic DNA from blood. Notably, while donors of the first two cohorts were almost exclusively Israeli Jews and Arabs, the WGBS samples were obtained from donors in the USA^17^, suggesting that this assay captures universal age-related methylation patterns.

### Performance of MageNet compared with existing epigenetic clocks

The original Horvath clock, using 353 individual CpGs measured using Illumina BeadChip arrays from whole blood, predicted age with a mean error rate of 3.9 years^20^. More recent clocks designed to predict chronological age reached accuracy down to 2.2 years when using 1000 CpGs^37^, while our own analysis of published array data resulted in accuracy of 1.89 to 2 years when using 30 to 80 CpGs^18^. The top performing algorithm described here, using deep targeted sequencing of two loci, ELOVL2 and C1orf132, combined with deep learning models, predicts chronological age with a median error of 1.7 years on unseen samples. Furthermore, we report an accuracy of 1.36 years on individuals under 50, and 0.9 years for individuals 35 or younger, representing a substantial improvement in accuracy. Notably, our model is based on a total of 17 CpG sites (nine at the ELOVL2 amplicon, and eight within C1orf132), but takes advantage of their binary combinatorial patterns. The superior performance of MAgeNet reflects the fact that it is robust to various environmental, clinical and hormonal changes that may affect biological, but not epigenetic, age prediction.

### Insights into the encoding of age by DNA

Our findings offer several insights into the biology of age encoding by DNA methylation. First, while the underlying biochemical mechanism remains a mystery, we found that age-related methylation changes occur across multiple adjacent CpGs, consistent with the typical regional nature of DNA methylation dynamics during development. Furthermore, we found that regional age-dependent methylation changes can occur either in a block-like coordinated manner, or independently and stochastically at each individual CpG site, as recently demonstrated^44,58^. This suggests that the molecular mechanisms underlying age-related methylation changes involve distinct pathways. We speculate that age-dependent activity of DNMT and TET enzymes is determined by factors such as DNA binding proteins, nucleosome positioning conferring steric hindrance, histone modifications and chromatin packaging. Further studies may look at these potential determinants at high resolution to understand how they dictate invariable methylation changes in specific loci. Most of the 45 markers we examined are gene-centered and overlap with regulatory regions, including promoters and genic regions, as well as CpG islands and polycomb CpG islands, as previously suggested for aging and cancer^7,59–61^, underscore the crucial role of DNA methylation in gene regulation, and highlights age-related effects on transcription programs. Second, the observation that deviations of clock prediction from chronological age are typically perpetuated to later measurements of the same individual, suggests that deviation is a one-time rare event; consequently, it appears that methylation changes at these loci are generally a faithful measure of elapsed time encoded in DNA, rather than a measure of chronological age in the formal sense. Future studies will address the time during which such deviations occur, and the potential determinants e.g. events taking place during differentiation and development, or genetic factors. We note that the lack of correlation between errors in age prediction and multiple environmental or physiological factors suggests that such early deviations are not a reflection of biological age.

Third, our findings reveal an interesting relationship to blood cell composition. We found that the best performing algorithm involved a locus that changed methylation with time (ELOVL2), and a locus that also reflected the characteristic age-related alterations in blood cell composition (C1orf132, marking T cells which are known to become less abundant in blood with advanced age)^49,62^. This also predicts that clocks using C1orf132 will be error-prone when an individual has altered blood counts e.g. during infection.

Fourth, the fact that model performance for individuals under 35 or 50 is better than the entire population (accuracy of 0.9 years, 1.36 years, or 1.7 years, respectively) suggests that in advanced age, remarkably small but nonetheless significant methylation noise accumulates which leads to reduced accuracy of encoding elapsed time.

Finally, our finding may shed some light on a fundamental question in the biology of aging - is biological age encoded by each individual cell or is it a function of a population of cells (e.g. the proportion of senescent cells). Our study does not answer this question, but it does show that at least elapsed time is encoded by a small number of cells, potentially in the methylation pattern of each cell, and can be accurately inferred from a small number of DNA molecules^63^.

### Practical implications

A straightforward utility of chronological age clocks is in the analysis of samples from unknown individuals, as often required in forensic case work. We note that the higher accuracy of our model for individuals under 35 or 50 years of age (test-set accuracy of 0.9 or 1.36 years, respectively) is beneficial in this regard, since most crime suspects are in this age range. We also note that the ability to infer chronological age from an extremely small number of cells is an important precondition for most forensic cases; for example, touch DNA typically allows the extraction of 0.5ng DNA, representing 100 genome equivalents. Finally, the current clock is optimized for blood DNA; forensic applications will require adaptation to additional body fluids such as saliva or sperm.

### Limitations and future directions

The methodology described here has several limitations, which present both challenges and opportunities for improvement. The reliance on PCR followed by sequencing introduces noise in the form of PCR duplicates, which likely accounts for much of our intra-assay variation. Duplicates can be avoided, for example by using unique molecular identifiers (UMIs), and we predict that scoring each template molecule only once will allow for a further increase in accuracy of the method. Another potential limitation of study is the narrow ethnicity of our donors – essentially just Jews and Arabs. However the ability of the model to infer age from WGBS of donors from the USA supports the generalizability of the model.

## Acknowledgments

We wish to thank Howard Cedar for insightful discussions, and members of the Dor and Kaplan labs for helpful discussions and comments. This work was supported by grants from the Israel Science Foundation grant (no. 1250/18, 259/23 to TK, 1065/16 to YD), from Horizon Europe (PANCAID consortium to TK and YD), to the Israeli Ministry of Science and Technology (A knowledge center for forensic DNA), and to the Center for Interdisciplinary Data Science Research at the Hebrew University. Research in the Dor lab is supported by the Helmsley Charitable Trust, and NCI (2U01CA210171-06). Yuval Dor holds the Walter and Greta Stiel Chair and Research grant in Heart studies. TK and YD are members of the Pamela and Paul Austin Research Center on Aging at the Hebrew University. The Jerusalem Perinatal Study was supported by NIH research grant no. R01HL088884, the Israel National Institute for Health Policy research grant (no. 2018/202). We thank Drs. Abed Nasereddin and Idit Shiff from the Interdepartmental Unit of the Hebrew University of Medicine for their support with DNA sequencing.

## Author contributions

YD, TK, BG, HH and RS conceived and designed this research. BLO, BG, and RS collected samples. BLO, AP, and SP performed experiments. DN, DC, NL, MV, RR and TK analyzed the data. IS, YF and HH provided JPS samples and helped with their interpretation. DN, YD, RS, and TK wrote the paper.

## Methods

### Sample collection

Population-based studies were approved by the ethics committee of Hadassah Medical Center. Procedures were performed under the Declaration of Helsinki. The donors have provided written informed consent.

### Library preparation

Blood samples were collected in EDTA tubes. DNA was extracted from 200 µl of blood using the “blood and tissue” Qiagen kit. Then, 500 ng of the solution was treated with bisulfite and amplified with PCR using primers designed for bisulfite-treated DNA. Pooled PCR products were subjected to multiplex NGS using the NextSeq 500/550v2 Reagent Kit (Illumina). Sequenced reads were separated by barcode, aligned to the target sequence on the human reference genome (hg19), and analyzed using custom programs as previously described^64^. Read pairs were merged, and sequenced DNA fragments were projected to a more compact methylation-specific representation, where non-CpG positions are discarded, methylated cytosines are denoted by C, and unmethylated ones by T, using wgbstools^65^.

### Non-linear age models using rate equations

Rate equations, and specifically ordinary differential equations (ODEs) describe the rate of change of a quantity with respect to time. The methylation dynamics of each CpG was modeled using two parameters, including the initial beta value (average methylation), as well as the relative rate of change, equivalent to the fixed percent of methylated molecules that undergo demethylation per year. These kinetics could be explicitly simulated using the Runge-Kutta method, or directly expressed using an exponentially decaying function. For CpG that gain methylation, the model assumes the percent of unmethylated CpGs that are methylated, per year. For the implementation and the derivation of the optimal rate of the ODE we used the *minimize* algorithm from the *scipy.optimize* library of Python 3.9, using L2 loss function.

### Conditioned Mutual Information

Pairwise mutual information was applied to quantify how coordinated or independent two CpGs are, while controlling for age-related changes in each CpG. Specifically, for each age ***k***, we estimated the pairwise mutual information ***I_k_(X;Y)*** was calculated for every pair of CpG sites ***X*** and ***Y***, by computing the difference between the marginal entropy ***H_k_(X)*** and the conditional entropy ***H_k_(X |Y)***. We then averaged across all ages, weighting by the number of samples available for each age. Entropies were calculated by merging all train-set samples from age ***k***, and using a Bayesian estimation of the average methylation per CpG. For this, we focused (for each age ***k***) on sequenced reads covering both CpGs, and counted the abundance of each binary combination (TT, TC, CT, CC) across the two CpGs ***X*** and ***Y***. The age-dependent probability of methylation ***P_k_(X=C)*** for CpG ***X*** was then estimated with a pseudocount of one, and the marginal entropy computed as minus the sum of ***P_k_(X=C) log_2_ P_k_(X=C)*** and ***P_k_(X=T) log_2_ P_k_(X=T)***. Similarly, the conditional entropy of ***X*** given ***Y*** was calculated as minus the sum of ***P_k_(X=a, Y=b) log_2_ P_k_(X=a, Y=b) / P_k_(Y=b)***, summing over four possible assignments for ***a*** and ***b***. High mutual information suggests that knowing the value of ***Y*** is informative of the value of ***X***, for each age ***k***. Further, as ***H_k_(X | X)*** equals zero, the self mutual information ***I_k_(X;X)*** equals the entropy of that CpG ***H_k_(X)***, in bits.

### Non-linear age models using generalized additive models (GAM)

Generalized additive models are a non-linear alternative to linear regression models. Here, the targeted variable is described as a linear sum of non-linear linkage functions applied to each predictor variable. For example, the age-dependent average methylation at each CpG site could be modeled using a spline or some smoothing function, which are then weighted and summed to predict age. The Python GAM implementation *pygam* (DOI 10.5281/zenodo.1208724) was used with default settings.

### Statistical tests

Effect of demographic and environmental traits, including sex, BMI, and smoking was tested using Python’s t-test implementation (scipy.stats.ttest_ind). Donors were binned by age, in 10-year intervals, and an equal number of donors were randomly sampled for each group. To compare the goodness-of-fit for different (nested) regression models, we used the F-statistic, which compares the relative improvements in fit (using the residual sum of squares), normalized for the number of sample and parameters: 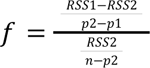, and p-value was assigned using the tail CDF of the F-distribution, and corrected for multiple hypothesis testing using Benjamini-Hochberg.

### Selection of age-correlated CpG sites

Prior to training the model, each CpG was tested independently for correlation with age using Spearman rank correlation, after grouping donors by age in a one-year bins. CpGs with Spearman correlation ≥ 0.8 that also showed a methylation range of change ≥ 20 percent points were selected for future analysis. For robustness, methylation range was defined by fitting a linear model to beta values and considering the absolute difference between predicted methylation at ages 20 and 80.

### Data processing and deep neural networks

Fragments were clipped to cover age-related CpGs (Table S3). Gapped fragments, or fragments with missing CpGs were ignored. The original 296 samples were split into train, test, and validation sets, stratified by age. Each sample was then augmented by generated 128 random subsets of 8,192 reads (sampled with replacement). For each set of fragments, covering a region of K age-related CpGs, three sets of methylation features were computed. First, we computed the average methylation level at each CpG (1 through K). Secondly, the abundance of fragments with exactly {0, 1, 2, …, K} methylated sites (out of K) was computed. Finally, the abundance of each of the 2^K^ possible methylation patterns, across the sequenced fragments was computed per sample. Features were concatenated to a single vector of length 2^K^+2K+1, serving as input for the network. A deepl learning architecture was designed based on fully connected neural networks, consisting of seven hidden layers of constant size. For regions of k≥4 CpGs, a hidden layer of 256 neurons were applied, or 512 neurons for *k*≥5, with a ReLU activation function. No regularization (pseudocounts) was applied for training. Different amplitudes of dropout were considered, as well as learning rate in the range of 3e-7 through to 3e-6. We used the ADAM optimization algorithm (with beta1 = 0.9, beta2 = 0.99, for weights update), with mini-batches of 128 samples. These parameters were selected using a grid search on the validation samples, using L1 loss. The deep learning model was implemented using the PyTorch library.

### Whole-genome bisulfite-sequencing data

Blood WGBS data was obtained from Loyfer et al.^17^. Data was analyzed using *wgbstools,* software suite we developed^65^, to convert the BAM files to binary DNA methylation data at single-fragment level (PAT files), and select fragments that fully cover the selected age-related CpGs, for each amplicon. Fragments were then augmented and processed as described for the PCR data.

### Model performance

Model performance on the test-set was assessed using the median absolute error (MAE), median absolute error for donors aged 50 or younger (MAE50), and root mean square error (RMSE and RMSE50). Each model was evaluated on the held-out test set. For each donor in the held-out test set, the sequenced data was sampled, with replacement, 128 times, age was predicted for each sample, and then averaged to produce a single chronological age prediction per sample.

## Supplementary Figures

**Supp. Figure 1:**
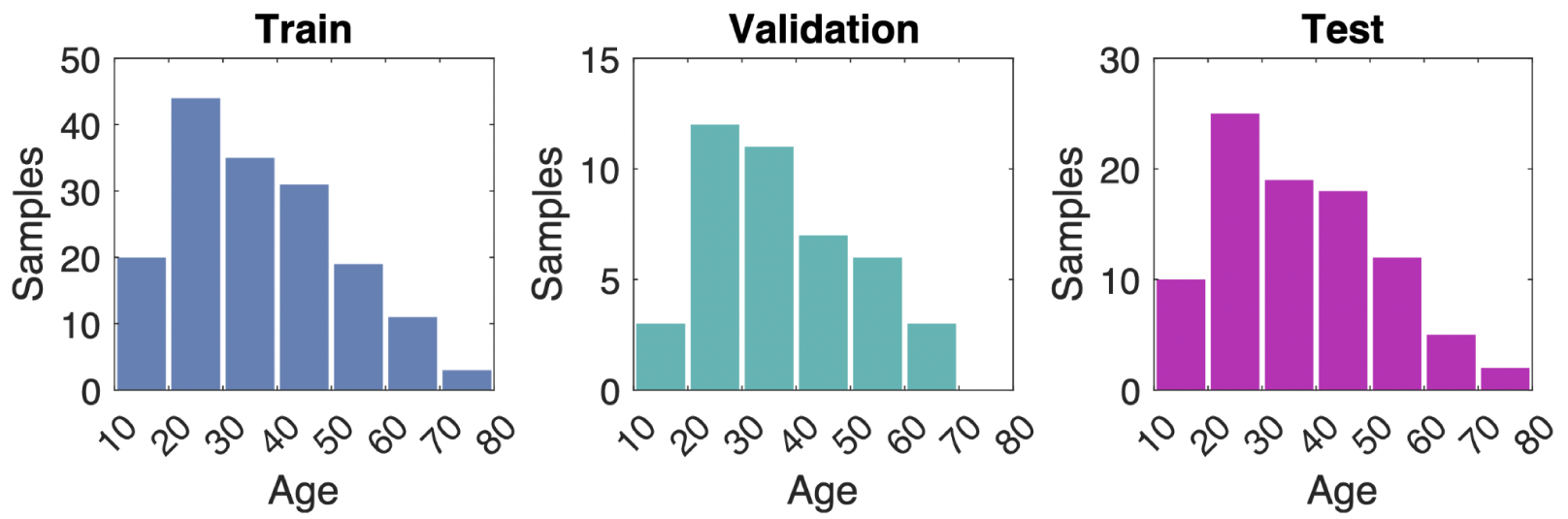
Age histogram for 205 training samples (left and middle) and 91 held-out test set samples (right). The train and test sets were split by a ratio of 70%/30% stratified by bins of 10 years, then the train set was split by a ratio of 80%/20% to train and validation sets.

**Supp. Figure 2:**
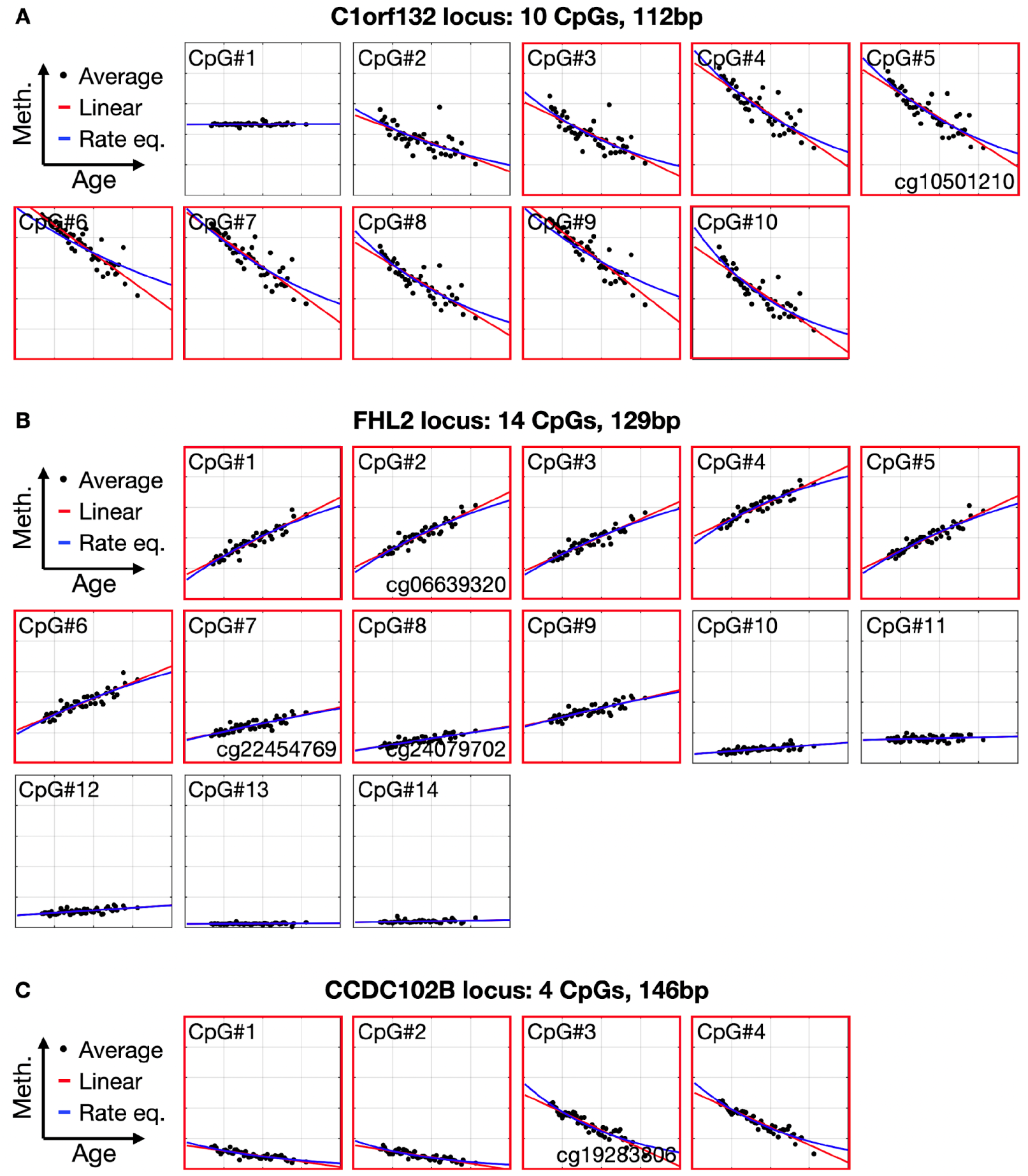
Same as. Figure 2**, for additional loci.** (**A)** C1orf132 (chr1:207996978-207997090, 10 CpGs), (**B**) FHL2 (chr2:106015715-106015844, 14 CpGs), and (**C**) CCDC102B (chr18:66389329-66389475, 4 CpGs).

**Supp. Figure 3:**
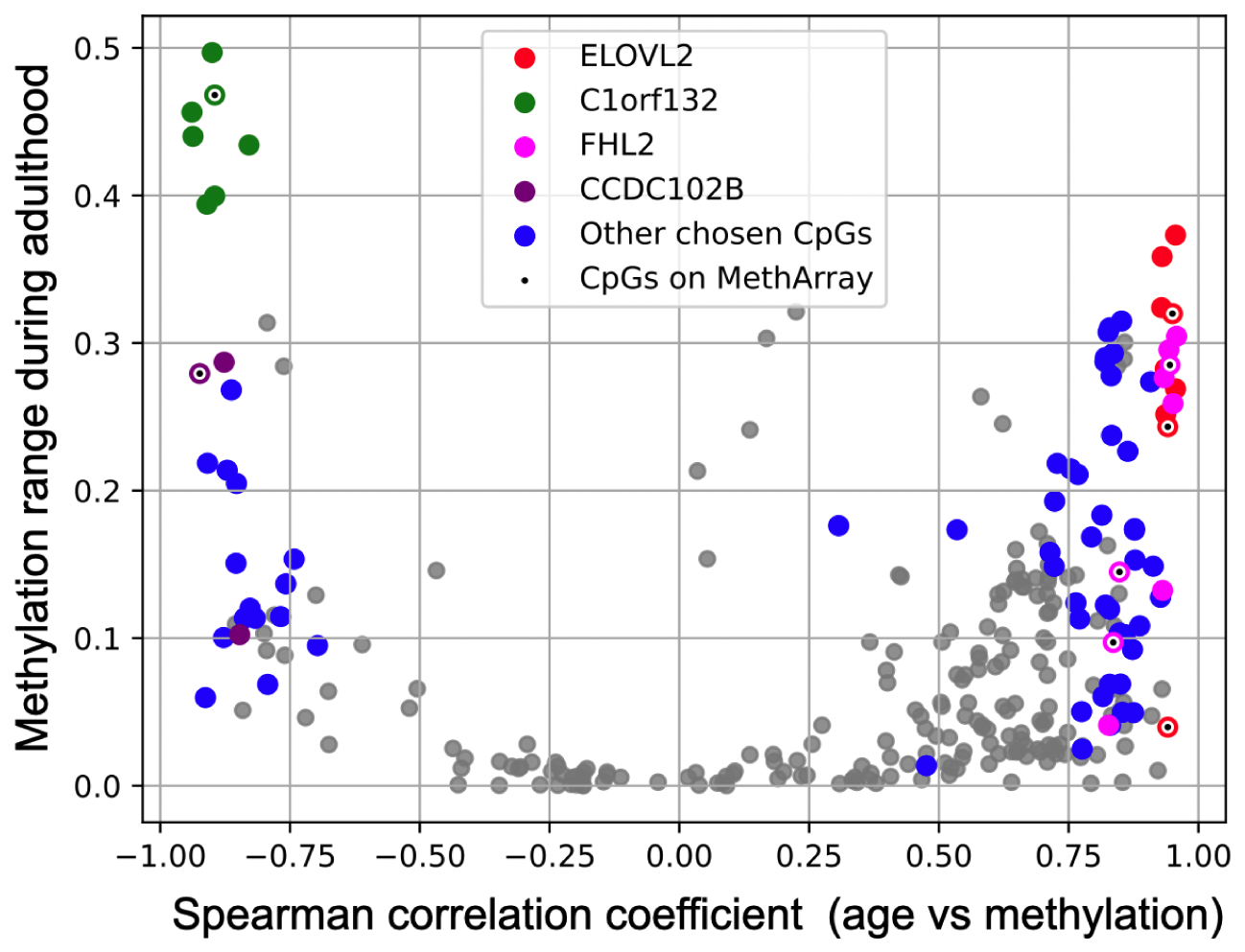
Clusters of age-related changes. For each amplicon (color) and for each CpG site (dot) we plot the Spearman correlation coefficient (X-axis) vs. the absolute range of methylation during adulthood (Y-axis). Intriguingly, CpGs that are the most correlated with age in each amplicon are not necessarily measured by 450K/EPIC arrays (black dot).

**Supp. Figure 4:**
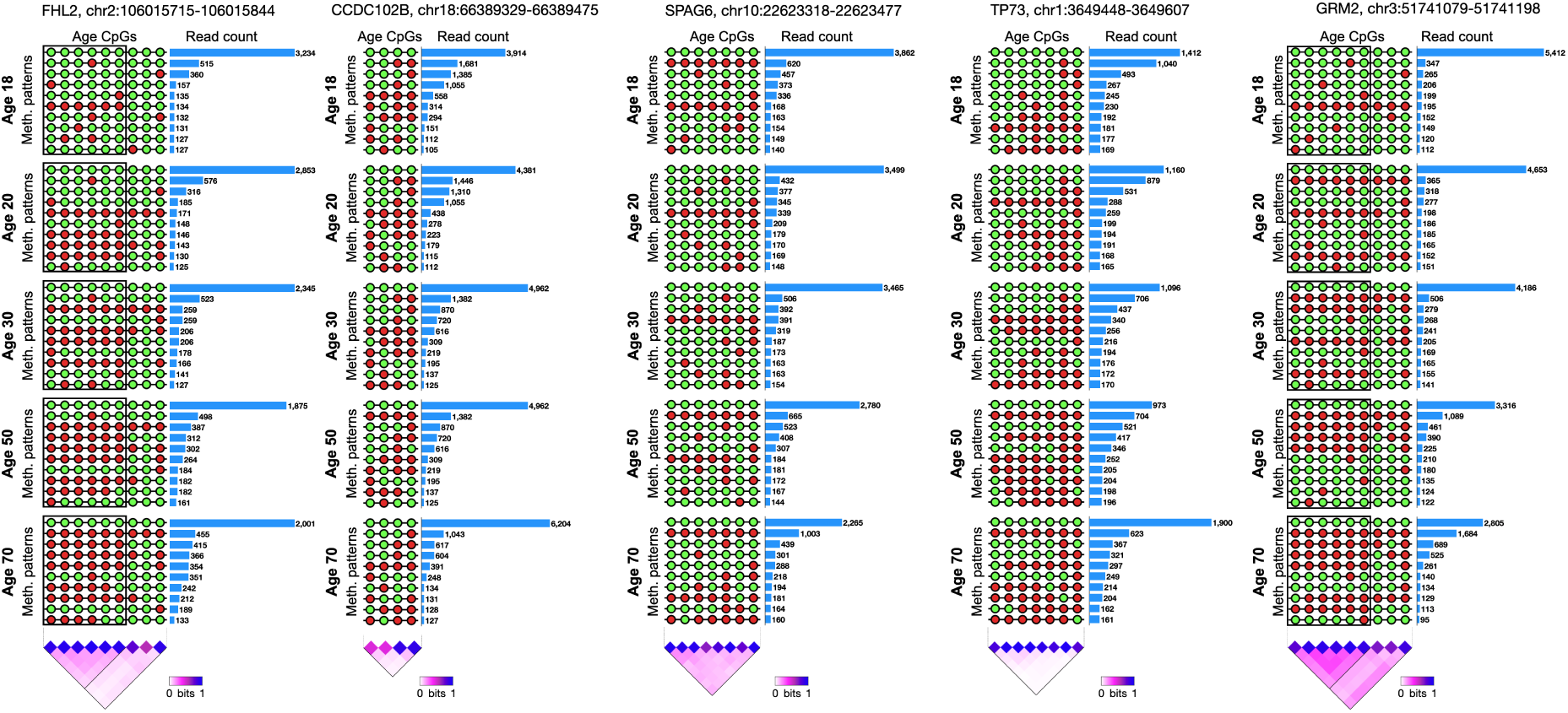
Same as Figure 3, for the FHL2, CCDC102B, SPAG6, TP73, and GRM2 locus. FHL2 shows block-like changes for the first six CpGs, out of nine age-related CpGs. GRM2 is highly coordinated.

**Supp. Figure 5:**
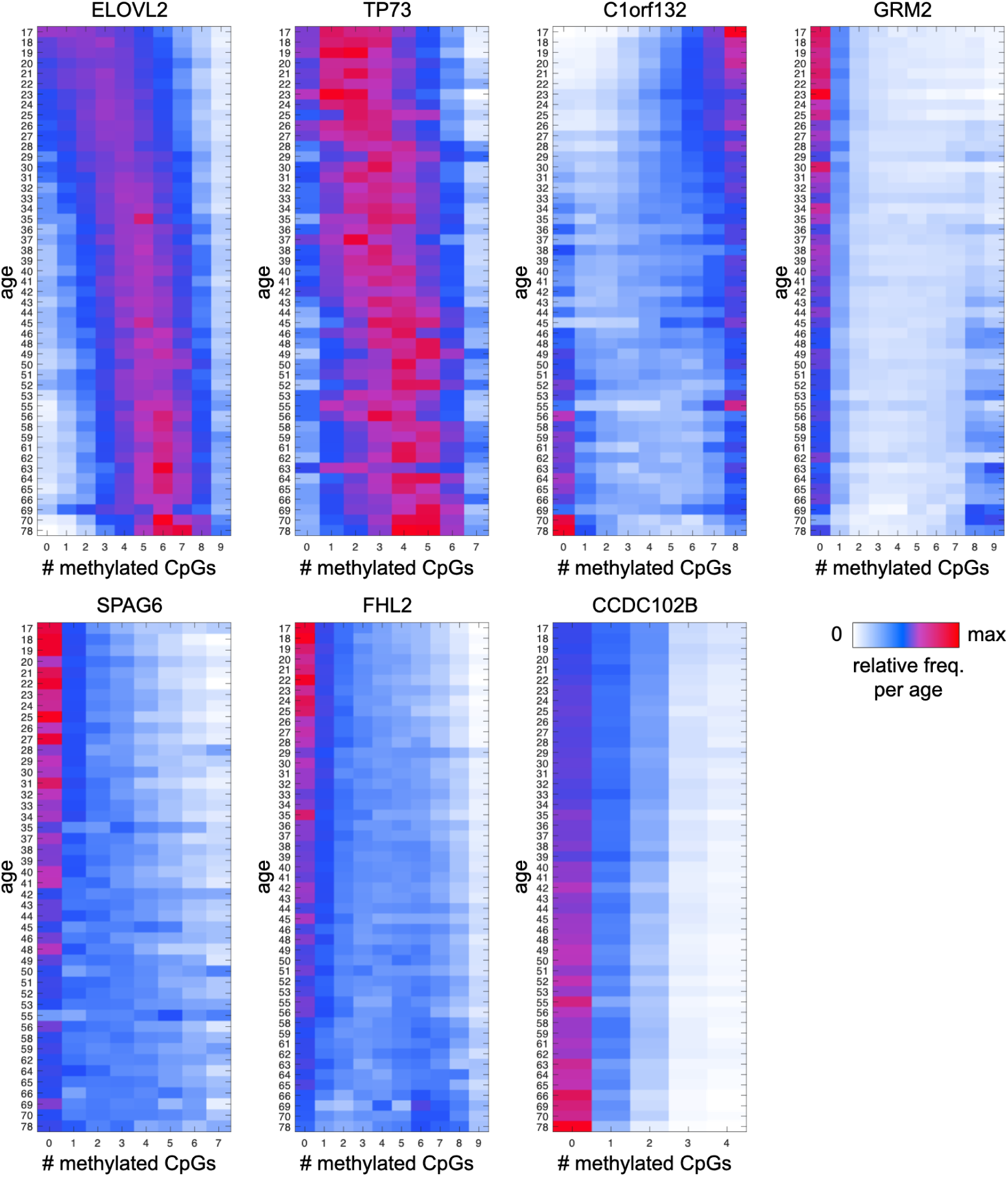
Stochastic gradual changes vs blocky (two-state) transitions. Each panel shows the prevalence of patterns with K methylated sites, from 0 (fully unmethylated, left) to all sites (fully methylated, right), for each sample, sorted by age (rows). ELOVL2 and TP73, for example, show a gradual change from mostly unmethylated fragments (young), to mostly methylated fragments (old). Conversely, C1orf132 shows a sharp blocky transition from fully methylated fragments (top right corner) to fully unmethylated ones (bottom right), whereas GRM2 shows a flipped blocky transition.

**Supp. Figure 6:**
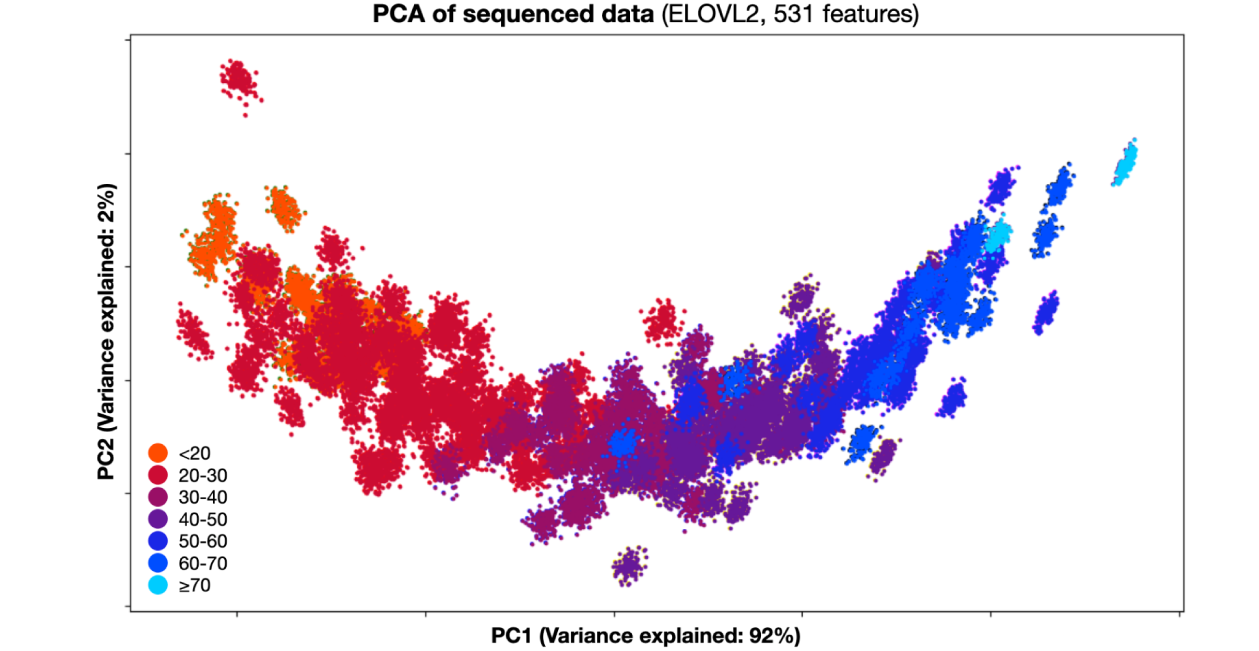
PCA analysis of the augmented high-dimensional data representation shows high concordance with age. Each cluster corresponds to one donor, where dots mark 128 randomly sampled subsets of 8192 reads. The high-dimensional representation of the data includes 531 features - 9 CpGs; 10 fragment-level features representing the abundance of fragments with exactly 0, 1, 2, through 10 methylated CpGs; and an additional 512 binary patterns (2 to the power of 9). Intriguingly, PC1 already captures 92% of the variance (X-axis), which is in agreement with chronological age.

**Supp. Figure 7:**
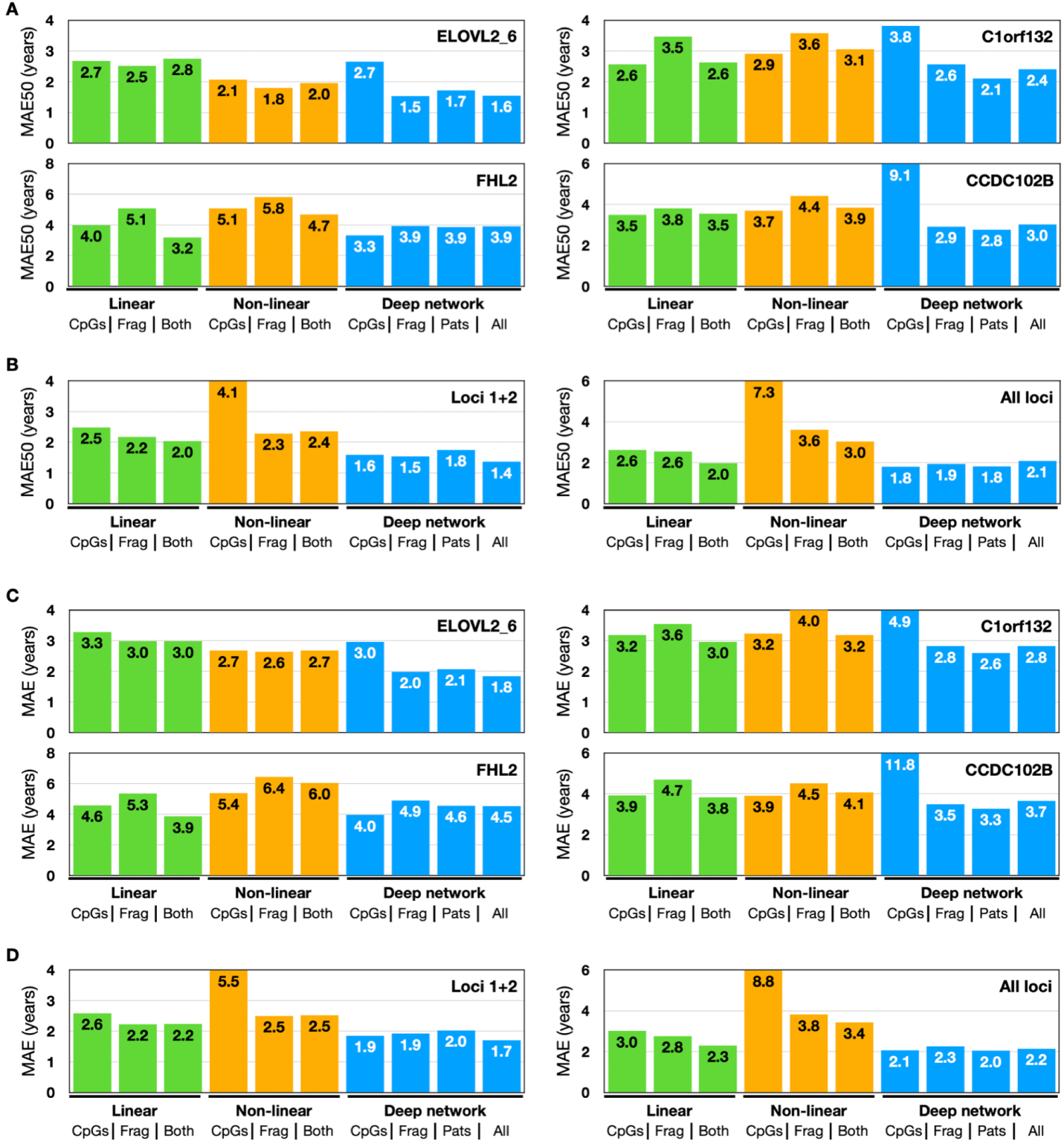
MAE50 and MAE across all trained models. Colors like in the main text: green for elastic-net, orange for GAM, and blue for neural network models. (A) MAE50 results of training the models for different input types, for each one of the four selected loci. (B) Same as A, but for the combination of loci, left for the combination of ELOVL2 and C1orf132, right for a combination of all four. (C) and (D) similar to A and B but the results are presented as MAE (includes all ages).

**Supp. Figure 8:**
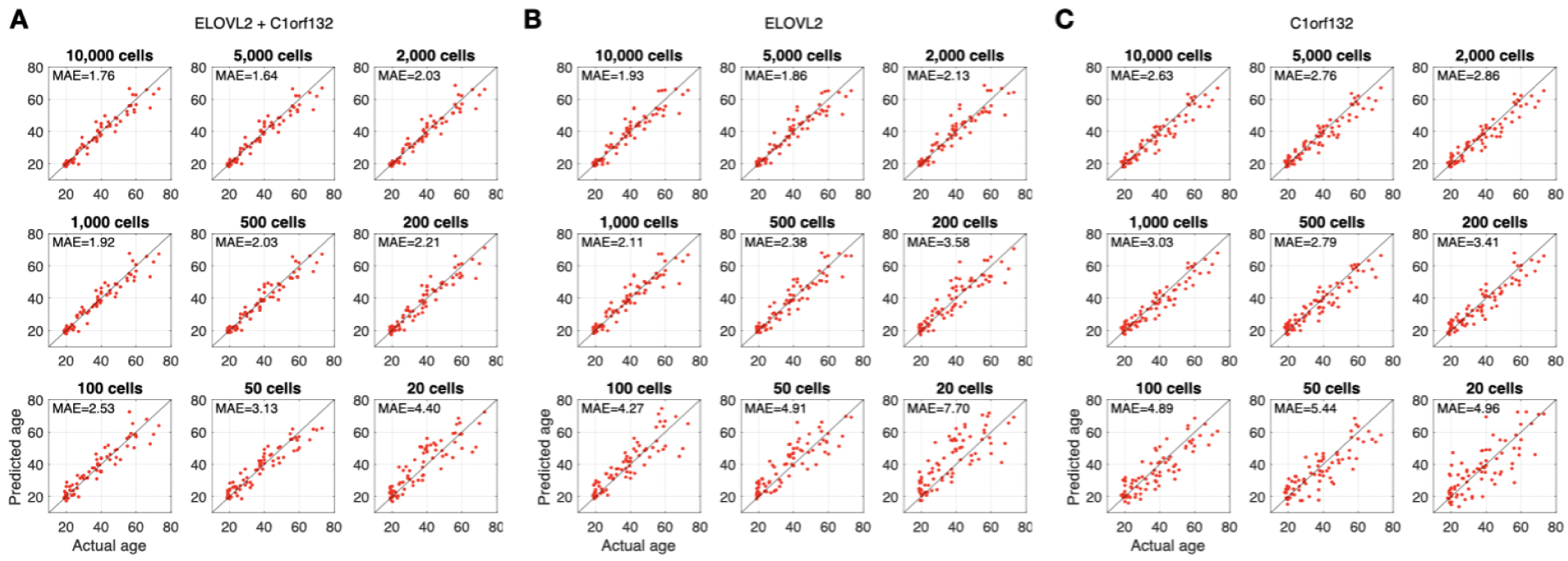
Amount of input material (in silico): The results of the simulation suggest that 500 cells may be enough for an accuracy similar to the regular model. Addition of strategies for removing duplicate reads may allow for accurate measurements even with a much smaller amount of starting material.

**Supp. Figure 9:**
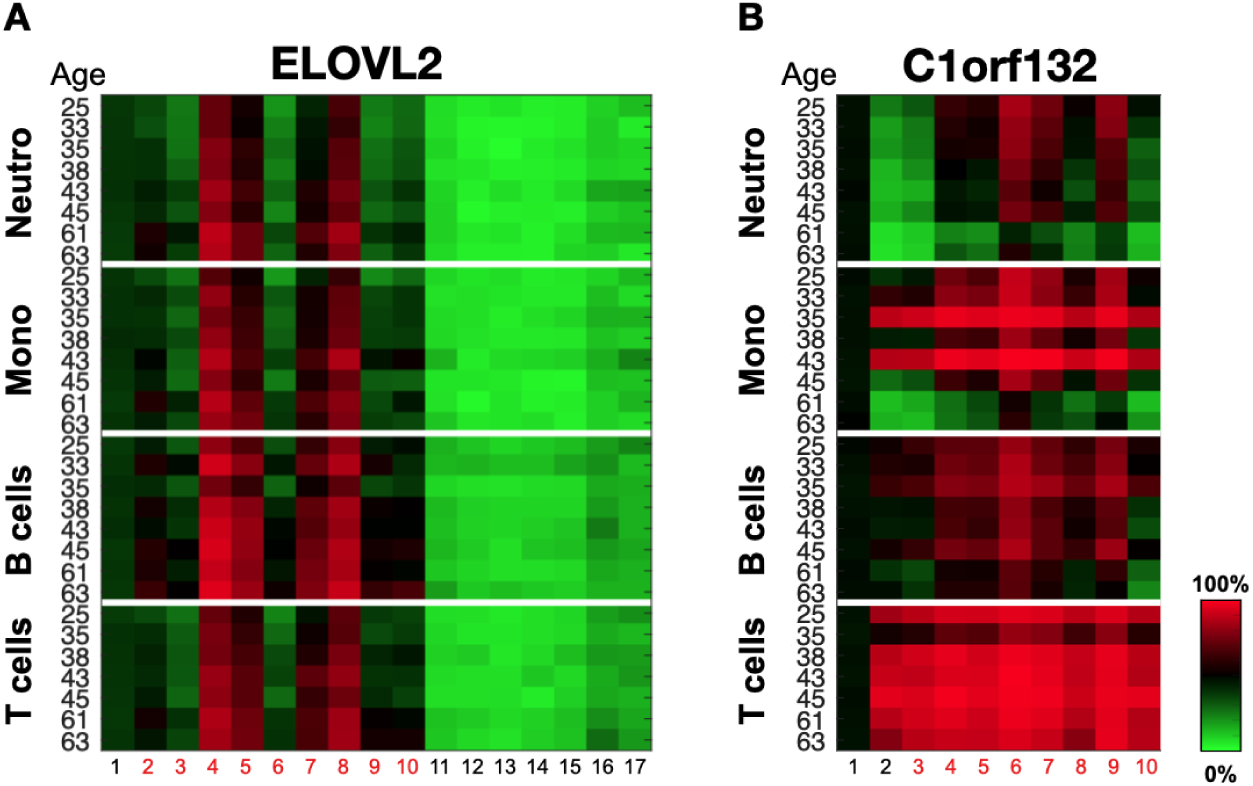
DNA methylation at purified blood cell types. (A) The ELOVL2 locus shows similar age-related methylation changes across various blood cell types, purified from donors aged 25 through 63. X-axis depicts individual CpGs. Other age-related markers (e.g. C1orf132) show some cell-type-specific effects, thus improving the model accuracy by integrating information on age-dependent blood cell composition.

**Supp. Figure 10:**
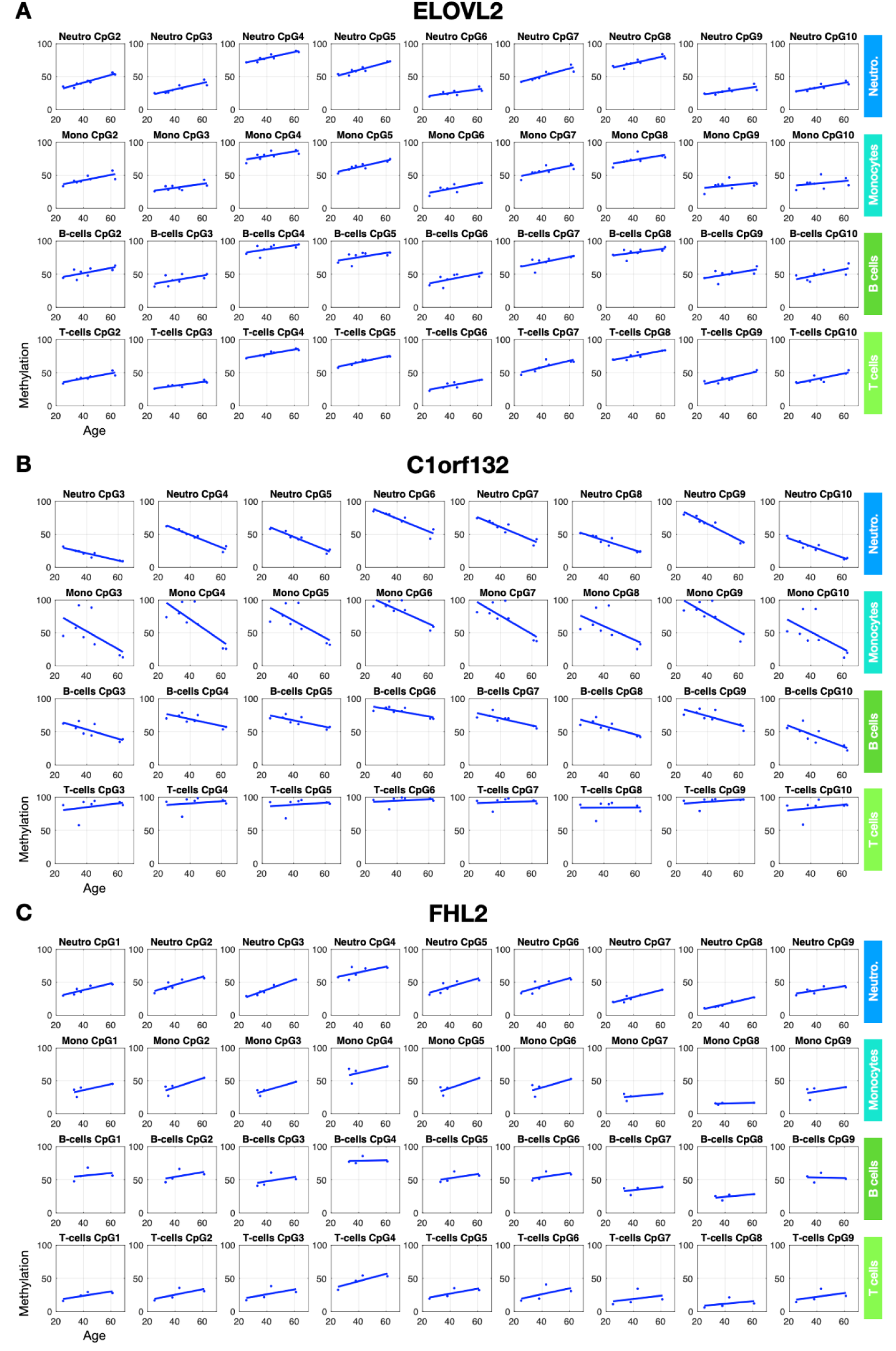
Age-related epigenetic changes for different loci and different blood cell types. **(A)** As for ELOVL2, all four tested cell-types show age-related methylation changes. **(B)** C1orf132 is a good example of the process of age-related demethylation. In this case, the age-related changes hardly affect the T-cell population, while other cell types lose methylation at different paces. **(C)** FHL2, an example of blockwise changes in age-relation methylation. All of the cell types present a slow pace of methylation gain.

**Supp. Figure 11:**
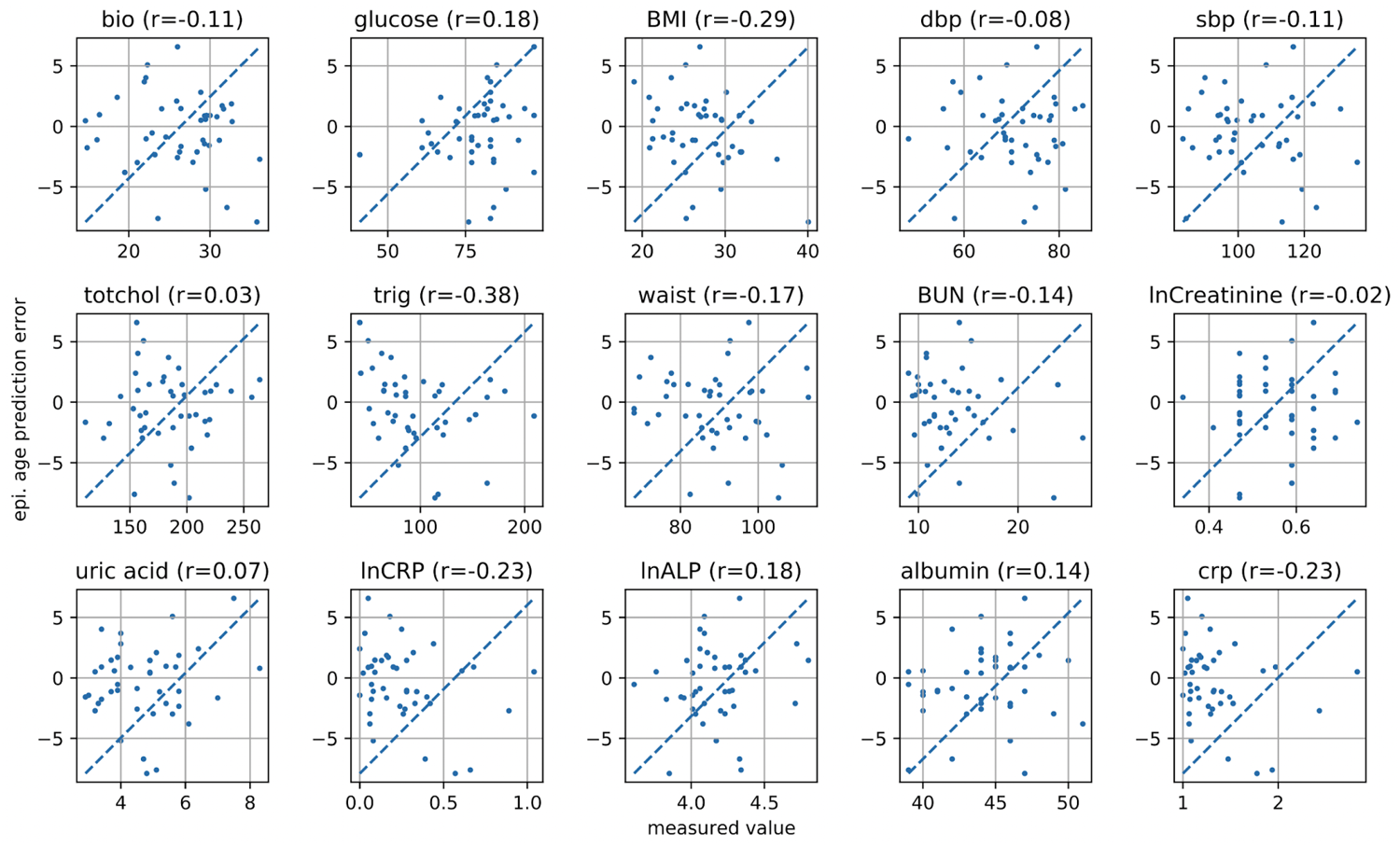
Bias in epigenetic age prediction is not correlated with biological age prediction or underlying measures. Epigenetic age prediction errors (top left) are not significantly correlated with biological age prediction errors, nor with other biological measures (except for triglycerides, r=-0.38).

## Supplementary Tables

**Supp. Table 1:**
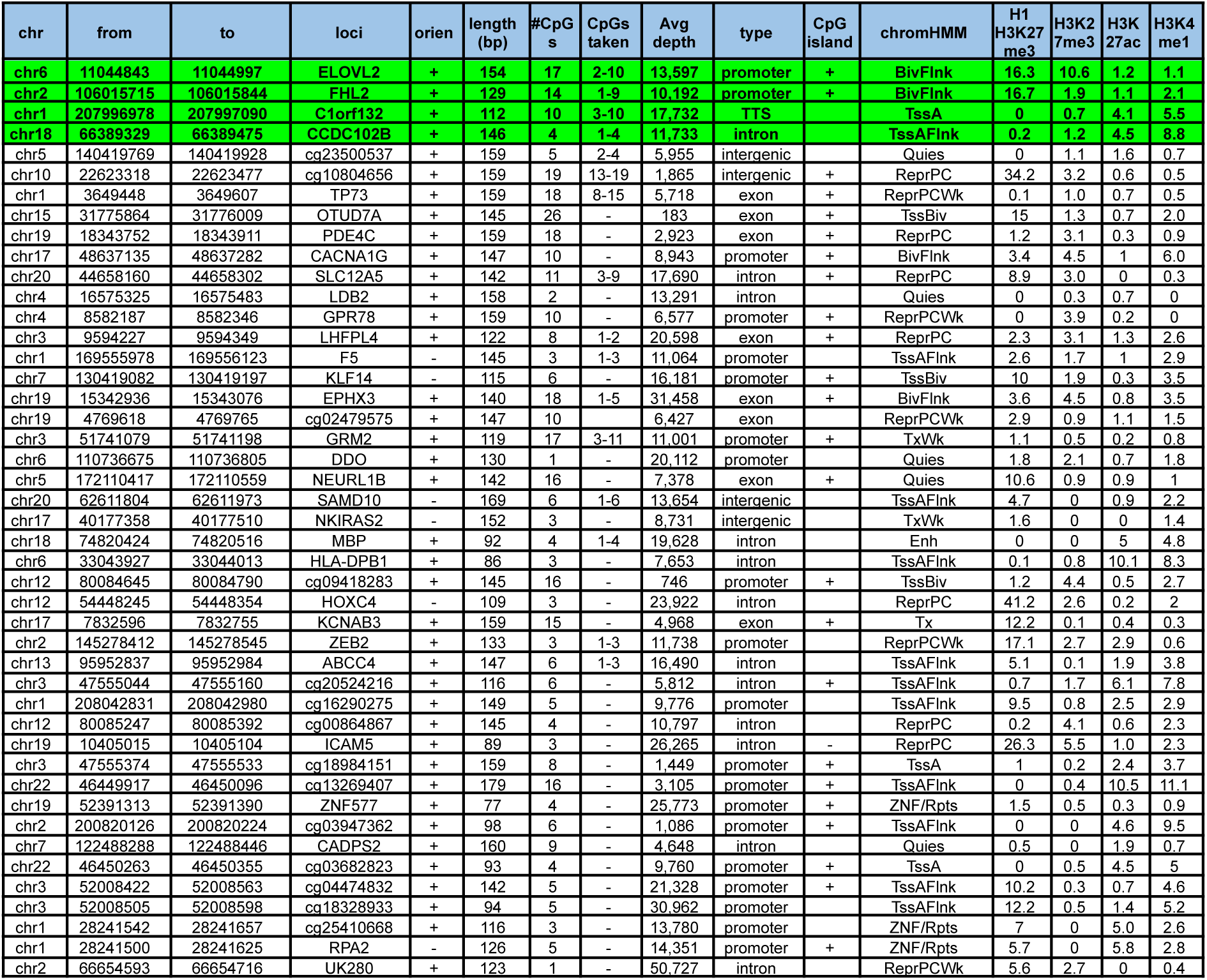
List of 45 genomic regions measured using targeted-PCR bisulfite sequencing (hg19). The top four age-predictive loci are highlighted in green. Information about the CpGs taken for the evaluation of the 16 loci is included. The targeted strand is always the top strand. chromHMM annotations are from the 15-state segmentation at PBMCs (E062). Chromatin ChIP-seq data is based on ENCODE data for H1 hESC and PBMCs (ENCFF832TSN, ENCFF150RIG, ENCFF759GIZ, ENCFF100NYH).

**Table S2** - Cohort demographics

**Supp. Table 3:** 16 age-associated genomic regions showing absolute Spearman’ based on core CpGs with correlation ≥ 0.8, with absolute change in methylation (ages 20-80) ≥ 20%.

|  | 1 | 2 | 3 | 4 | 5 | 6 | 7 | 8 | 9 | 10 | 11 | 12 | 13 | 14 | 15 | 16 | 17 | 18 | 19 |
| --- | --- | --- | --- | --- | --- | --- | --- | --- | --- | --- | --- | --- | --- | --- | --- | --- | --- | --- | --- |
| TP73 | 0.04 | 0.75 | 0.73 | 0.72 | 0.76 | 0.78 | 0.73 | 0.79 | 0.81 | 0.82 | 0.82 | 0.78 | 0.81 | 0.36 | 0.83 | 0.77 | 0.74 | 0.79 |  |
| F5 | -0.90 | -0.91 | -0.91 |  |  |  |  |  |  |  |  |  |  |  |  |  |  |  |  |
| C1orf132 | 0.19 | -0.81 | -0.85 | -0.91 | -0.90 | -0.95 | -0.93 | -0.91 | -0.95 | -0.92 |  |  |  |  |  |  |  |  |  |
| FHL2 | 0.95 | 0.95 | 0.95 | 0.94 | 0.96 | 0.94 | 0.85 | 0.85 | 0.84 | 0.65 | 0.21 | 0.62 | 0.17 | 0.25 |  |  |  |  |  |
| ZEB2 | -0.85 | -0.86 | -0.86 |  |  |  |  |  |  |  |  |  |  |  |  |  |  |  |  |
| LHFPL4 | 0.90 | 0.90 | 0.61 | 0.73 | 0.54 | 0.65 | 0.65 | 0.73 |  |  |  |  |  |  |  |  |  |  |  |
| GRM2 | 0.04 | 0.10 | 0.89 | 0.88 | 0.88 | 0.89 | 0.88 | 0.87 | 0.87 | 0.87 | 0.87 | 0.88 | 0.87 | 0.88 | 0.89 | 0.88 | -0.12 |  |  |
| cg23500537 | 0.57 | 0.90 | 0.93 | 0.86 | 0.77 |  |  |  |  |  |  |  |  |  |  |  |  |  |  |
| ELOVL2 | 0.24 | 0.94 | 0.95 | 0.95 | 0.95 | 0.96 | 0.96 | 0.97 | 0.95 | 0.95 | 0.78 | 0.77 | 0.66 | 0.73 | 0.79 | 0.74 | 0.51 |  |  |
| cg10804656 | 0.73 | 0.70 | 0.74 | 0.79 | 0.82 | 0.83 | 0.85 | 0.78 | 0.73 | 0.62 | 0.72 | 0.73 | 0.76 | 0.74 | 0.78 | 0.83 | 0.89 | 0.78 | 0.79 |
| ABCC4 | -0.88 | -0.87 | -0.88 | -0.79 | -0.81 | -0.85 |  |  |  |  |  |  |  |  |  |  |  |  |  |
| CCDC102B | -0.88 | -0.91 | -0.97 | -0.97 |  |  |  |  |  |  |  |  |  |  |  |  |  |  |  |
| MBP | -0.86 | -0.89 | -0.91 | -0.88 |  |  |  |  |  |  |  |  |  |  |  |  |  |  |  |
| EPHX3 | 0.89 | 0.91 | 0.90 | 0.87 | 0.87 | 0.87 | 0.77 | 0.82 | 0.86 | 0.69 | 0.80 | 0.54 | 0.74 | 0.74 | 0.75 | 0.59 | 0.59 | 0.52 |  |
| SLC12A5 | 0.56 | 0.55 | 0.82 | 0.90 | 0.88 | 0.89 | 0.88 | 0.83 | 0.84 | 0.63 | 0.62 |  |  |  |  |  |  |  |  |
| SAMD10 | -0.77 | -0.74 | -0.72 | -0.85 | -0.84 | -0.82 |  |  |  |  |  |  |  |  |  |  |  |  |  |

**Supp. Table 4:**
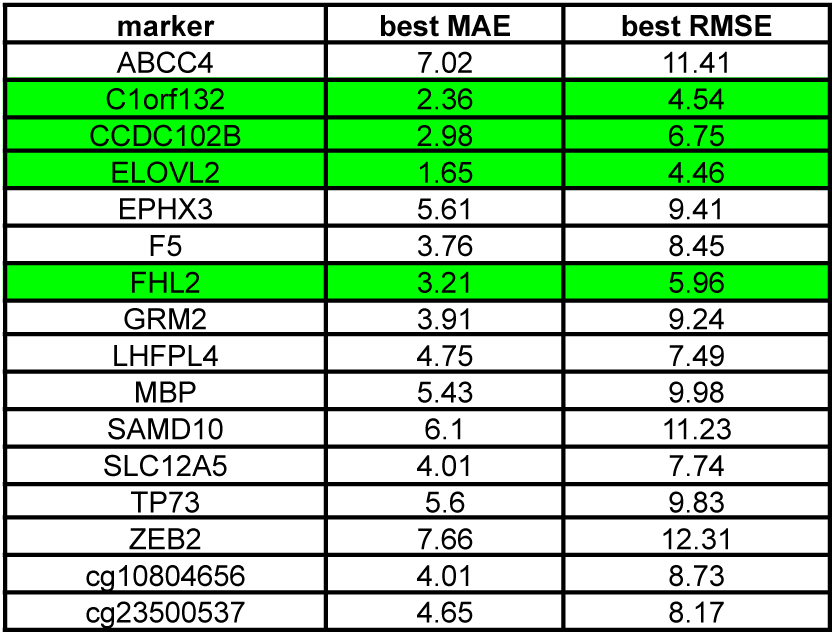
Age-related markers and their best MAE and RMSE values. Green marks MAE≤4 and RMSE<7.

**Supp. Table 5:** MAE50 of the three types of model for the different types of information.

|  | ElasticNet<br>CpGs | ElasticNet<br>frags | ElasticNet<br>CpGs+frags | GAM<br>CpGs | GAM<br>frags | GAM<br>CpGs+frags | MAgeNet<br>CpGs | MAgeNet<br>frags | MAgeNet<br>patterns | MAgeNet<br>full |
| --- | --- | --- | --- | --- | --- | --- | --- | --- | --- | --- |
| ELOVL2 | 2.67 | 2.52 | 2.75 | 2.07 | 1.80 | 1.96 | 2.65 | 1.54 | 1.72 | 1.55 |
| C1orf132 | 2.57 | 3.47 | 2.62 | 2.91 | 3.58 | 3.06 | 3.82 | 2.56 | 2.11 | 2.40 |
| FHL2 | 3.98 | 5.07 | 3.19 | 5.08 | 5.81 | 4.67 | 3.31 | 3.94 | 3.87 | 3.91 |
| CCDC102B | 3.49 | 3.80 | 3.54 | 3.70 | 4.42 | 3.85 | 9.09 | 2.92 | 2.76 | 3.02 |
| All 4 | 2.63 | 2.55 | 1.97 | 7.32 | 3.61 | 3.03 | 1.80 | 1.94 | 1.83 | 2.09 |
| 1+2+3 | 2.40 | 2.23 | 1.57 | 7.70 | 3.15 | 2.81 | 1.90 | 1.59 | 1.43 | 1.88 |
| 1+2 | 2.48 | 2.17 | 2.04 | 4.07 | 2.29 | 2.36 | 1.59 | 1.54 | 1.75 | 1.36 |
| 1+2+4 | 2.71 | 2.74 | 2.16 | 5.75 | 3.34 | 2.49 | 1.54 | 1.93 | 1.83 | 1.75 |

**Supp. Table 6:**
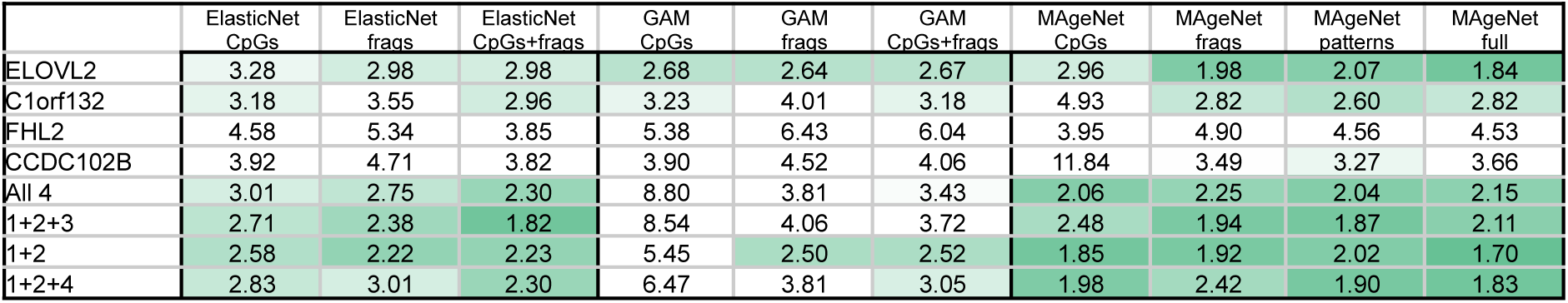
MAE of the three types of model for the different types of information.

|  | ElasticNet<br>CpGs | ElasticNet<br>frags | ElasticNet<br>CpGs+frags | GAM<br>CpGs | GAM<br>frags | GAM<br>CpGs+frags | MAgeNet<br>CpGs | MAgeNet<br>frags | MAgeNet<br>patterns | MAgeNet<br>full |
| --- | --- | --- | --- | --- | --- | --- | --- | --- | --- | --- |
| ELOVL2 | 3.28 | 2.98 | 2.98 | 2.68 | 2.64 | 2.67 | 2.96 | 1.98 | 2.07 | 1.84 |
| C1orf132 | 3.18 | 3.55 | 2.96 | 3.23 | 4.01 | 3.18 | 4.93 | 2.82 | 2.60 | 2.82 |
| FHL2 | 4.58 | 5.34 | 3.85 | 5.38 | 6.43 | 6.04 | 3.95 | 4.90 | 4.56 | 4.53 |
| CCDC102B | 3.92 | 4.71 | 3.82 | 3.90 | 4.52 | 4.06 | 11.84 | 3.49 | 3.27 | 3.66 |
| All 4 | 3.01 | 2.75 | 2.30 | 8.80 | 3.81 | 3.43 | 2.06 | 2.25 | 2.04 | 2.15 |
| 1+2+3 | 2.71 | 2.38 | 1.82 | 8.54 | 4.06 | 3.72 | 2.48 | 1.94 | 1.87 | 2.11 |
| 1+2 | 2.58 | 2.22 | 2.23 | 5.45 | 2.50 | 2.52 | 1.85 | 1.92 | 2.02 | 1.70 |
| 1+2+4 | 2.83 | 3.01 | 2.30 | 6.47 | 3.81 | 3.05 | 1.98 | 2.42 | 1.90 | 1.83 |

**Supp. Table 7:** Adjusted p-values (FDR-corrected) for sex, BMI, and smoking effects. Age is shown as a positive control.

|  | ELOVL2 | C1orf132 | FHL2 | CCDC102B | 1+2 | 1+2+3 | 1+2+4 | All loci |
| --- | --- | --- | --- | --- | --- | --- | --- | --- |
| BMI (18.5-24.9) vs. < 18.5 or > 24.9 | 0.89 | 1 | 0.87 | 0.45 | 0.83 | 0.74 | 0.89 | 1 |
| Female vs. Male | 1 | 0.33 | 0.79 | 0.79 | 0.91 | 0.68 | 0.81 | 0.91 |
| Never vs. current smoker | 0.25 | 0.79 | 0.87 | 0.79 | 0.96 | 0.79 | 0.79 | 0.4 |
| Former vs. Current smoker | 0.63 | 0.4 | 0.85 | 0.25 | 0.79 | 0.4 | 0.4 | 0.4 |
| Former vs. Never-smoked | 0.79 | 1 | 0.68 | 1 | 0.81 | 0.79 | 0.85 | 0.89 |
| Smoking years <5 vs >10 | 0.96 | 0.68 | 0.79 | 0.96 | 0.79 | 0.79 | 0.66 | 0.79 |

**Supp. Table 8:** Spearman rho and adjusted p-values (FDR-corrected) for biological age features effects on MAgeNet predicted age deviations from chronological age.

|  | Spearman rho | p-value | FDR |
| --- | --- | --- | --- |
| Biological age | -0.09 | 0.36412 | 0.39538 |
| Glucose | 0.17 | 0.09165 | 0.16586 |
| BMI | -0.23 | 0.01981 | 0.10755 |
| dbp | -0.09 | 0.38674 | 0.38176 |
| sbp | -0.1 | 0.29686 | 0.35816 |
| totchol | 0.02 | 0.8357 | 0.60496 |
| trig | -0.31 | 0.0016 | 0.017374 |
| waist | -0.11 | 0.2533 | 0.34381 |
| BUN | -0.05 | 0.63496 | 0.53036 |
| lnCreatinine | -0.04 | 0.72074 | 0.55901 |
| uric acid | 0.08 | 0.41798 | 0.37822 |
| lnCRP | -0.22 | 0.02617 | 0.094722 |
| lnALP | 0.18 | 0.06057 | 0.13154 |
| albumin | 0.13 | 0.19359 | 0.3003 |
| crp | -0.22 | 0.02707 | 0.073485 |

